# Automated high-throughput selection of DNA aptamers using a common optical next-generation sequencer

**DOI:** 10.1101/2024.06.24.600375

**Authors:** Alissa Drees, Christian Ahlers, Timothy Kehrer, Natascha Ehmke, Alice Frederike Rosa Grün, Charlotte Uetrecht, Zoya Ignatova, Udo Schumacher, Markus Fischer

**Author notes:** Yusuf Hamied Department of Chemistry, University of Cambridge, Lensfield Road, Cambridge CB2 1EW, United Kingdom (Alissa Drees). Corresponding author: Prof. Dr. Markus Fischer Hamburg School of Food Science Grindelallee 117, 20146 Hamburg.

## Abstract

Aptamers are conventionally selected via ‘Systematic Evolution of Ligands by Exponential Enrichment’ (SELEX). This process is, however, laborious, time-consuming, and has a relatively low efficacy. Here, we developed an automated high-throughput screening platform for the selection of DNA aptamers which consists of an optical next-generation sequencer with a modified software and hardware to automatically perform fluorescence-based binding assays on the displayed DNA sequences subsequent to sequencing. Using this platform, after only three to five SELEX rounds we selected highly affine DNA aptamers for the lectins LecA and LecB of *Pseudomonas aeruginosa* as well as for the *Pseudomonas* Exotoxin A. In comparison, twelve rounds of conventional SELEX resulted in three-fold less affine aptamers for LecA and PEA and none for LecB. Our high throughput-approach bears great potential to augment SELEX as it significantly increases time efficiency, enabling the selection of aptamers within only one week.

---

Aptamers are synthetic single-stranded nucleic acid oligomers that are able to bind affinely to target structures. The sequence of these oligonucleotides determines their structural conformation and thus their interactions with the target. While aptamers can achieve a similar affinity and specificity as antibodies, they offer several advantages such as higher thermal stability, *in vitro* selection and synthesis, which allows for higher batch stability, low synthesis costs, and the possibility of straightforward modifications^1, 2^. Hence, aptamers have great potential as receptors in detection systems and for therapeutic purposes^1, 2^.

The selection of aptamers was first described in 1990^3, 4^. Conventionally, aptamer selection is performed by the iterative ‘Systematic Evolution of Ligands by Exponential Enrichment’ (SELEX) process. Common to all SELEX variants is that the selection itself is difficult to monitor and control and, with about 20–30%, has a relatively low success rate^5, 6^. In addition, the SELEX process is usually quite laborious and time-consuming.

A more recent method with enormous potential for the selection of aptamers is ‘High-Throughput Sequencing Fluorescent–Ligand Interaction Profiling’ (HiTS-FLIP)^7^. It enables both high-throughput screening and quantitative measurement of the affinity of DNA-protein interactions. For this approach, HiTS-FLIP exploits the potential of flow cells of optical sequencers to display millions of DNA sequences on which fluorescence-based affinity and specificity assays can be performed directly after sequencing. This methodology increases significantly the selection efficiency of aptamers. In addition, it allows for addressing previously unanswerable questions about the selection process (e. g. the influence of various selection conditions on the affinity enrichment) and the relationship between sequence, structure, and function of DNA-protein interactions.^8^

In this study, we demonstrate the use of a common, commercially available sequencer – the Illumina MiSeq – as a straightforward, automated platform for high-throughput measurement of ssDNA-protein interactions.

With this approach, we successfully selected and characterised DNA aptamers for three proteins of *Pseudomonas aeruginosa*, i. e. the two lectins LecA and LecB as well as the *Pseudomonas* Exotoxin A (PEA). These three proteins are rather difficult targets for aptamer selection due to their relatively low isoelectric points (theoretical pIs: LecB: 3.88, LecA: 4.94 and PEA: 5.36)^9^. By comparing directly to conventional aptamer selection via SELEX, we achieved the selection of more than three-fold more affine aptamers for LecA and PEA and for the first time the selection of aptamers for LecB, for which none were obtained via SELEX. In addition, our method reduced the required time for aptamer selection from several months to one week. Our data show that HiTS-FLIP is a powerful tool for aptamer selection. Furthermore, the approach bears great potential for various other biochemical questions such as the determination of the sequence specificity and affinity of transcription factors^7^ and of Cas-enzymes^10^ in nucleotide resolution.

## Results

### Automated high-throughput aptamer array for customised binding assays

We worked on two options on how to modify an Illumina MiSeq sequencer to perform automated fluorescence-based quantitative binding measurements on the DNA clusters immobilised on the flow cell:

The first option enables completely automated HiTS-FLIP experiments, but includes a minor hardware modification, similarly as described for the N2A2 array^11^. Briefly, an external valve was connected to the originally vacant port 23 of the internal valve of the MiSeq so that additional reagents can be selected and directed to the flow cell during the experiment (Fig. S1). We synchronised the external valve by encoding its desired position via specific ‘waiting times’ – not used proprietarily by Illumina – in the recipe. Upon recording of those ‘waiting times’ in a log file as the recipe was executed, they were recognised by a customised Python script, which then triggered a switch of the external valve.

The second option allows to perform HiTS-FLIP experiments without requiring any hardware modification. For this approach, a custom-filled reagent cartridge replaced Illumina’s proprietary reagent cartridge before the binding assays. In comparison to the first option, this approach enables the constant cooling of all reagents, including the protein dilutions, and requires less amount of the reagents, since the lines do not have to be re-primed for each use. A manual exchange of the cartridges subsequent to sequencing was required, thus only a semi-automation of the experiment was achieved using this option.

For both options the software of the sequencer was modified according to the requirements of the experiment to be performed. In specific, the steps included in the ‘second read’ were changed to a restriction enzyme digestion followed by multiple cycles of washing steps (i. e. with 100% formamide and 50 mM NaOH), an incubation of the flow cell with fluorescently labelled target, and imaging.

For the HiTS-FLIP experiments, we used aptamer libraries pre-enriched by few SELEX rounds (i. e. 3 to 5), which reduced their diversity to enable a more comprehensive display of the remaining sequences on a MiSeq’s flow cell^12, 13^. The aptamer libraries (Fig 1A) were sequenced on the modified MiSeq and after the paired-end turnaround converted into functional aptamers (Fig. 1B). For this, the 3’ constant region was cleaved off by EcoRI. The degree of EcoRI restriction was monitored by hybridisation of a fluorescently labelled oligonucleotide complementary to the 3’ constant region of the aptamer library prior to and after restriction (Extended Data, Fig. 4). During a subsequent heating step, the 5’ constant region of the library was hybridised to a complementary oligonucleotide to reduce its interference with the folding of the random region^14^. After generation of the aptamer display, the binding assays using fluorescently labelled target proteins were performed as shown in Fig. 1C. For this procedure, the flow cell was incubated with different concentrations of an AF647-labelled protein (i. e. PEA, LecA, or LecB) and imaged after a washing step. To ensure good focussing and alignment of these images, besides phiX, also a so-called fiducial mark (FM) oligonucleotide was spiked into the aptamer library. During the binding assay, fluorescently labelled oligonucleotides complementary to phiX’s adapters and to the FM oligonucleotide were then hybridised to the clusters displayed on the flow cell. This approach enabled optical focussing during the imaging of the binding assay as well as alignment of the images^11^. The FM’s mean fluorescence intensities were also used to check the required comparability of the determined fluorescence intensities of different cycles. Binding parameters, such as the equilibrium dissociation constant *K*_d_, could be determined for all displayed clusters using the captured images. For this procedure, first, the fluorescence intensities on channel A of each cluster at a given cycle had to be extracted, as automatically performed by the MiSeq’s real-time analysis (RTA). Secondly, by plotting the fluorescence intensities against the target concentration for each cluster, binding functions, such as a HILL function, were fitted, from which the *K*_d_ values were determined^15^.

**Figure 1:**
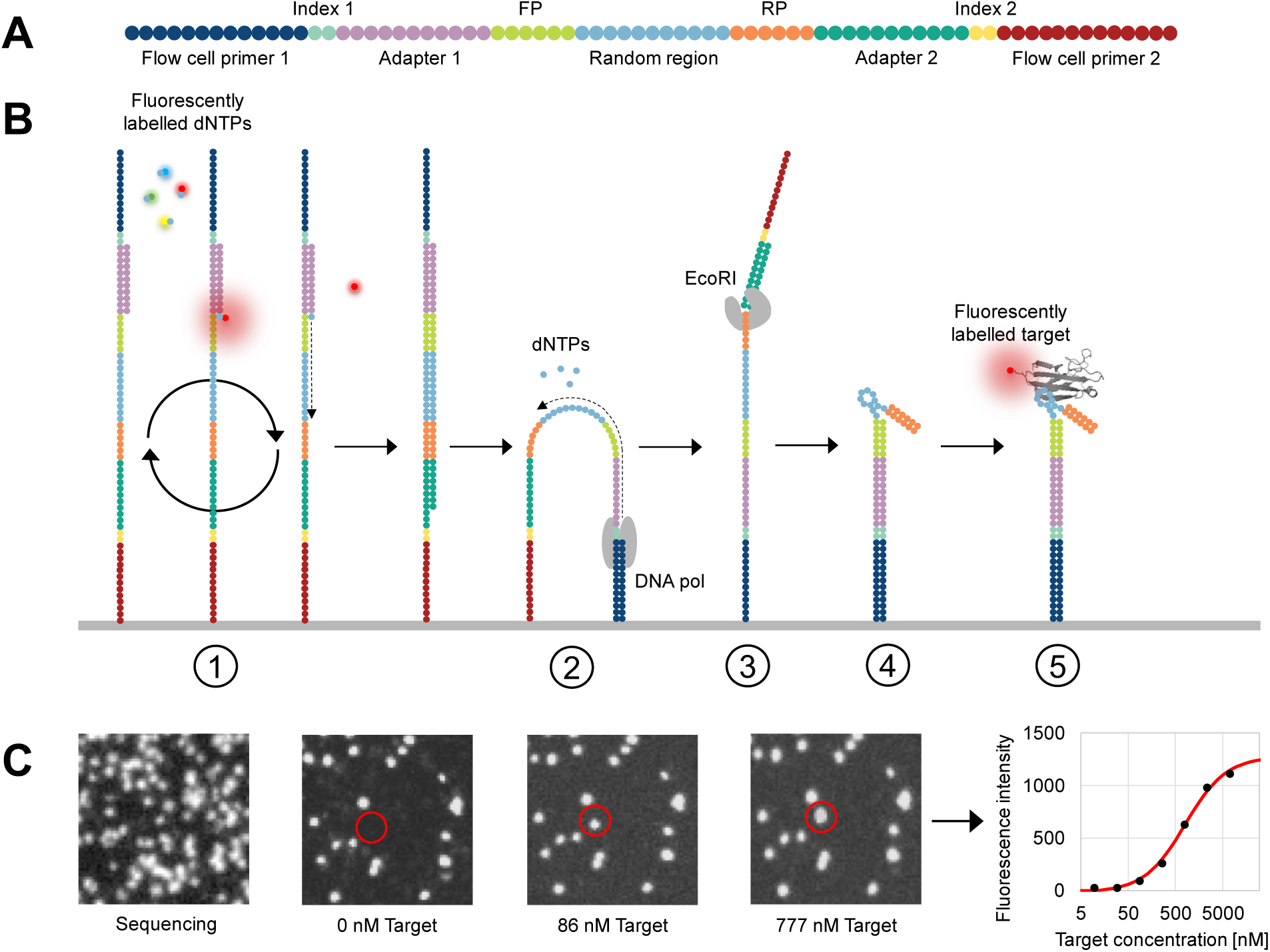
The principle of aptamer selection via HiTS-FLIP. (**A**) The used libraries’ structural composition. (**B**) The experiment consists of five steps shown schematically for a single strand immobilised on the flow cell: Sequencing-by-Synthesis (1), a paired-end turnaround (2), cleavage of the 3’ constant regions (3), hybridising of complementary oligonucleotides to the remaining constant regions and folding of the aptamers (4), and the binding assay performed by incubation with fluorescently labelled target followed by washing and imaging of the flow cell (5). (**C**) The flow cell with images of channel A acquired during sequencing (left), as well as during the binding assay with different concentrations of AF647-labelled PEA. By plotting the fluorescence intensity extracted from the images against the concentration of PEA, binding curves can be fitted, from which parameters such as the *K*_d_ can be determined – for each cluster on the flow cell.

In order to perform reliable binding assays using an optical sequencer, it has to be taken into account that the MiSeq is not designed to have an accurate absolute determination of fluorescence intensities, but rather enables the determination of relative differences between the four fluorescence channels^16^. On the one hand, this is due to the instrument’s optics facilitating a centred illumination of the flow cell’s tiles. On the other hand, the determined fluorescence intensities are depending on the clusters’ size, which can differ due to amplification bias during the cluster generation^17^. Therefore, it is important to consider each cluster’s own maximum fluorescence in the fit of binding curves.

### Selection of aptamers for PEA

Aptamers for PEA were selected by HiTS-FLIP using an N_50_ DNA aptamer library pre-enriched by three SELEX rounds. We chose this pool which exhibited 1.5–10 million unique sequences as quantified by next-generation sequencing (NGS). The highest enriched sequences had a relative count lower than 0.05%. In contrast, after another SELEX cycle the diversity of the aptamer pool decreased to approx. 250 thousand unique sequences with the most enriched sequence reaching up to 2.73%. Consequently, the PEA3 pool was estimated to enable the display of a greater diversity of sequences on the MiSeq’s flow cell, than subsequent pools, while already providing a certain enrichment of sequences.

Subsequent to sequencing and folding of the displayed ssDNA sequences, we introduced PEA labelled with AF647 to the flow cell in seven different concentrations ranging from 422 pM to 6.91 µM.

Over 450 thousand clusters displaying sequences of the aptamer library passed the applied quality and length filters (max. 3 bases under Q30 and 48–52 nt after primer trimming). Most of those clusters did not show any binding towards PEA, which can be considered a successful negative control for the experiment (Fig. 2A). By fitting a HILL-function for each cluster, we uncovered multiple suitable aptamer candidates. In total, six potential aptamers were selected for further characterisation (Fig. 2B). We considered clusters with diverse cluster sizes and shapes, different maximum fluorescence intensities and varying proximity to FM clusters. Of the six sequences, five bound with high affinity (< 1 µM) to PEA as determined by a surface plasmon resonance spectroscopy (SPR) assay. Only PEA194456 did not show binding interactions with PEA, which might be caused by the 3’ immobilisation of the aptamers during the SPR assay. For PEA7290, PEA33764, and PEA170478 the *K*_d_ values (150– 300 nM) measured by HiTS-FLIP could be confirmed by SPR as well as by affinity chromatography (Table 1). In addition, we confirmed the binding also in solution by native nano-electrospray ionisation mass spectrometry (nano-ESI-nMS, see supplemental material). Multiple aptamers with an over 25-fold increased affinity in comparison to the only aptamer previously published for PEA – PEA R14.33 (*K*_d_ = 4.2– 4.5 µM), selected after 14 rounds of SELEX^18^ – were obtained by HiTS-FLIP. This result emphasises the potential of HiTS-FLIP in selecting high-affinity aptamers.

**Figure 2:**
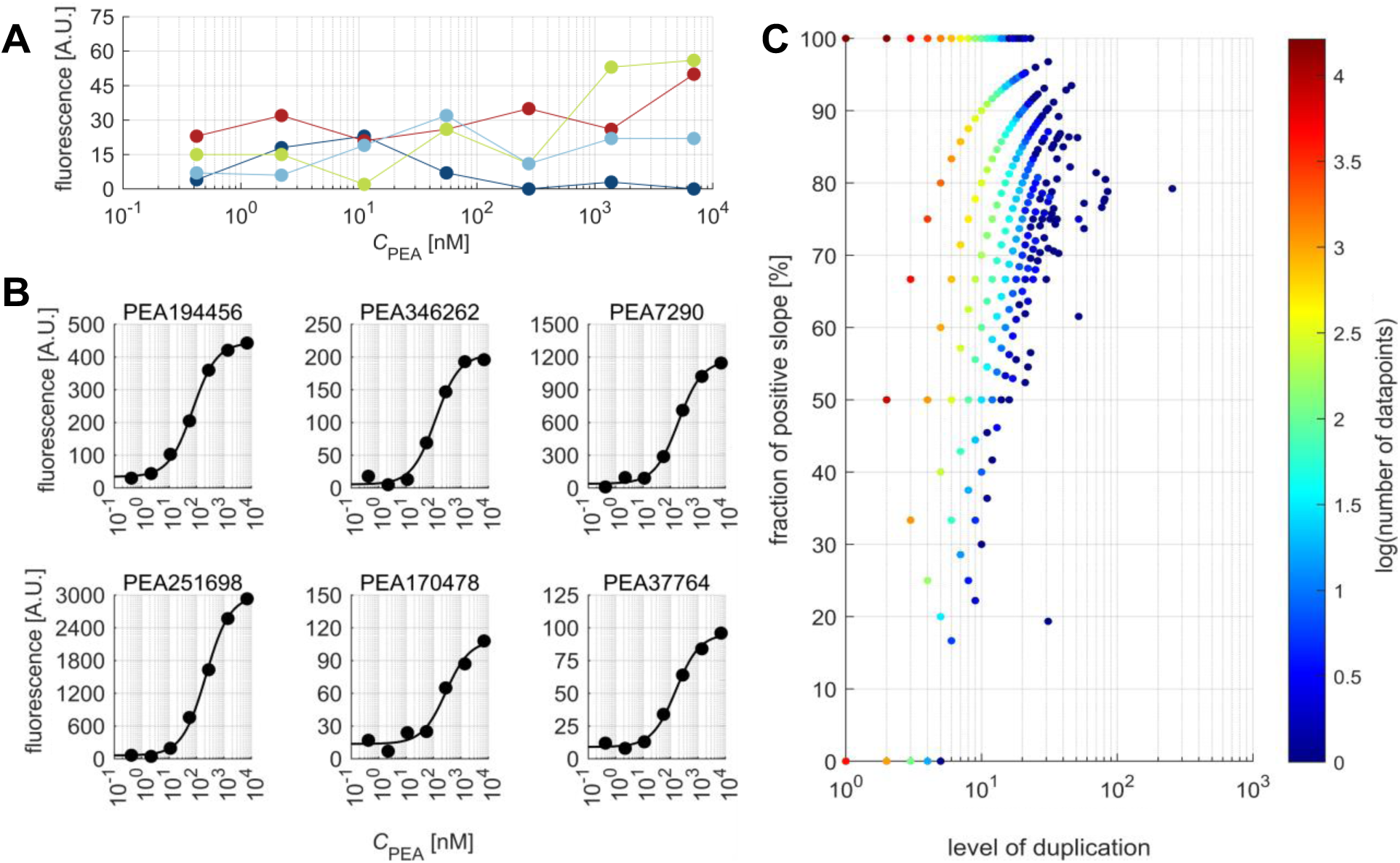
Results of the HiTS-FLIP for PEA. (**A**) Four examples of the fluorescence intensities of non-binding clusters plotted against the PEA concentration. (**B**) Scatter plots and HILL-fits of selected clusters. (**C**) Relationship between sequence frequency and increase of fluorescence intensity of PEA: This scatter plot illustrates the proportion of sequences exhibiting a positive slope between the mean of the first and last three intensities based on their frequency within the flow cell. The colour bar indicates the number of sequences which constitute to each point in the scatter plot.

**Table 1:**
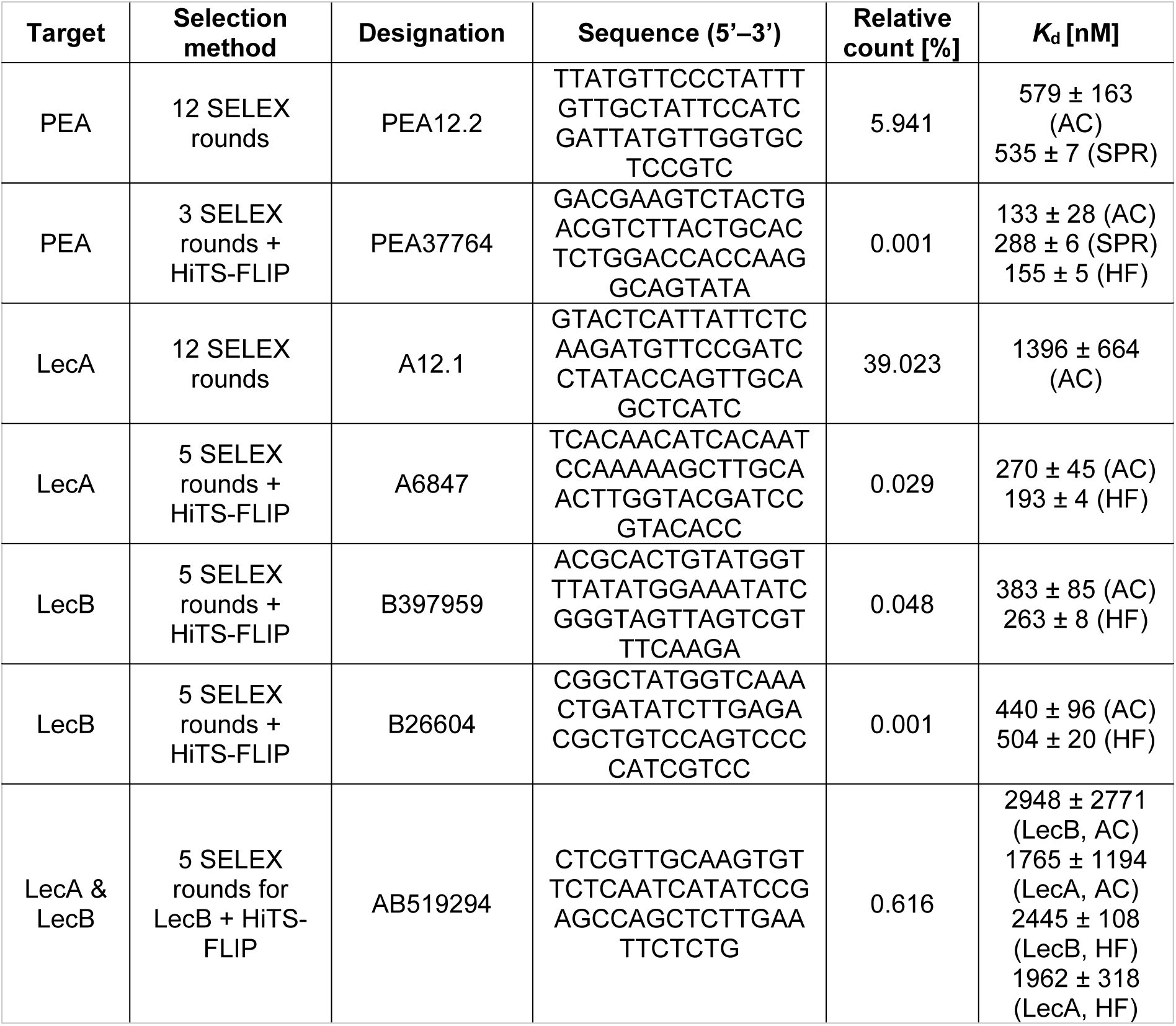
Summary of the best selected aptamers for LecA, LecB, and PEA via SELEX and HiTS-FLIP. AC: Affinity chromatography; HF: HiTS-FLIP.

HiTS-FLIP can also provide more comprehensive insights into the interactions between the target and the displayed sequences^19–21^. *Inter alia*, the success of previously applied enrichment strategies can be determined^22^. For the HiTS-FLIP for PEA, a positive correlation between the abundance of a sequence and its affinity, as described by an increase of the measured fluorescence value from the lowest to the highest protein concentration, was observed (Fig. 2C). This finding shows that the previous three SELEX rounds already successfully enriched affine sequences. However, after twelve rounds of SELEX only three-fold less affine aptamers were among the ten most abundant sequences. In detail, the most affine aptamer selected by SELEX was PEA12.2, whose affinity was determined to be 535 ± 7 nM by SPR and 596 ± 219 nM by affinity chromatography (Table 1).

### Selection of aptamers for LecA and LecB

To determine whether HiTS-FLIP could be used to select aptamers specific for one or two targets simultaneously, aptamers for the two calcium-dependent, tetrameric lectins of *Pseudomonas aeruginosa* – LecA and LecB^23^ – were selected by HiTS-FLIP using a library combining two pools pre-enriched by five SELEX rounds for either target. By NGS, these pools of aptamers for LecA and LecB were determined to have a diversity of approx. 600 and 200 thousand sequences, with the highest enriched sequences having a share of up to 0.1% and up to 4%, respectively.

In total, 136,095 clusters of LecA5 and 213,960 LecB5 clusters passed the applied quality and length filters. During the binding assay, the influence of different buffers (i. e. PBST or Mg- and Ca-containing TBST) was tested. The three technical replicates of each of the ten target concentrations (ranging from 11.4 nM to 5.83 µM for AF647-LecA and from 13.0 nM to 6.66 µM for AF647-LecB) in TBST revealed a high reproducibility of the measured fluorescence intensities with a mean coefficient of variation of 9.25% (mean standard deviation of 12 fluorescence units, both at highest target concentration as well as during blanks).

We were able to select monospecific aptamers with affinities of up to 200–300 nM for either lectin and bispecific aptamers with an affinity in the low micromolar range for both lectins (Fig. 3), as validated by affinity chromatography (Table 1). Interestingly, the aptamer B397959 showed high affinity towards LecB in both analysed buffers, while B26604 only bound to LecB in the Ca- and Mg-containing TBST buffer, suggesting a Ca-dependence of the interaction of B26604 with the lectin. Since the sugar-binding of the two lectins is Ca-dependent^23^, we reasoned that a Ca-dependency might be an indicator of the inhibitory effect of the selected aptamers, that would make the aptamers interesting for therapeutic applications^24, 25^. Thus, to analyse the inhibitory properties of the LecB-binding aptamers, we performed immunohistochemistry using human nasal concha tissue. The likely Ca-dependent aptamer B26604 displayed a stronger inhibition of LecB binding compared to the other tested aptamers (Fig. 3). However, in comparison to the L-fucose control, the inhibitory effect was not strong enough to be of clinical relevance (IC_50_ > 50 µM). Nonetheless, these DNA aptamers for LecB are, to the best of our knowledge, the first aptamers successfully selected for a protein with an isoelectric point of less than four (LecB’s theoretical pI: 3.88)^9^.

**Figure 3:**
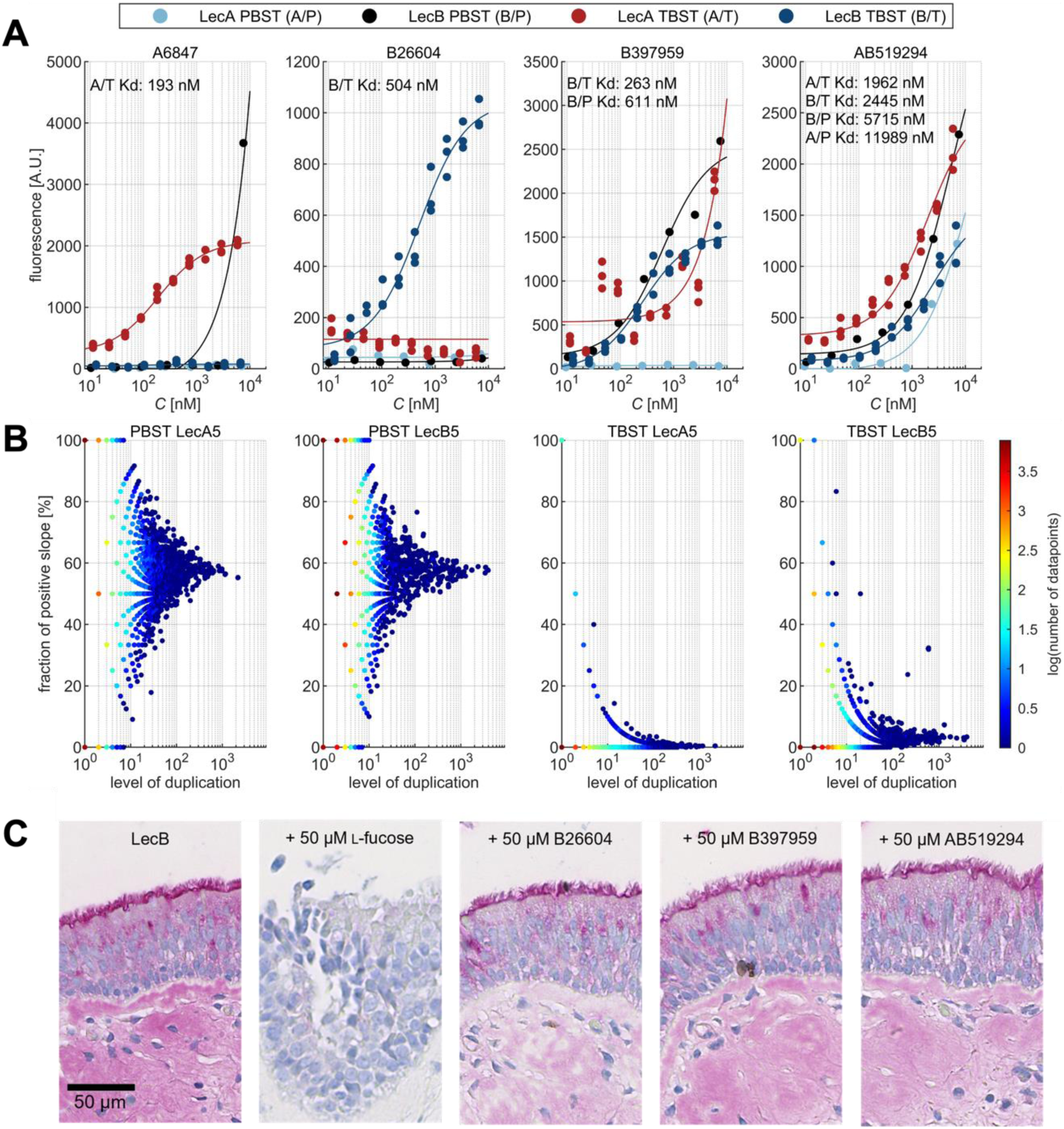
Results of the HiTS-FLIP experiment for LecA and LecB. (**A**) Scatter plots and HILL-fits of selected clusters’ fluorescence during the incubation with LecA and LecB in PBST or TBST. The outlying fluorescence value for A6847 at the highest concentration for LecB (> 3500) in PBST are most likely to be attributed to an integer error during background subtraction by the MiSeq’s RTA or saturated pixels during image acquisition. (**B**) Relationship between sequence frequency and increase of fluorescence intensity of LecA and LecB in PBST and TBST: These scatter plots illustrate the proportion of sequences exhibiting a positive slope between the mean of the first and last three intensities based on their frequency within the flow cell. The colour bars indicate the number of sequences which constitute to each point in the scatter plot. (**C**) Analysis of inhibitory properties of L-fucose and selected aptamers on LecB’s binding to sections of human nasal concha tissue by lectin immunohistochemistry showing that L-fucose leads to a complete inhibition of LecB’s binding to the surface epithelium and the underlying connective tissue, while the aptamers only reduce the binding.

In contrast, after twelve selection rounds of conventional SELEX, for LecA only aptamers with affinities ≥ 1.3 µM (Table 1) were selected. For LecB no aptamers were obtained after twelve SELEX rounds, which showcases a low selection efficiency, as also observed by HiTS-FLIP (Fig. 3B). This finding emphasizes the importance of the insight that HiTS-FLIP can provide by enabling the detection of lower abundance affine sequences in a diverse pool of sequences.

## Discussion

The idea to extend the scope of optical next-generation sequencers to perform ultra-high-throughput binding assays has been described already a decade ago^7^, however, only recently the suitability of the Illumina MiSeq to perform automated binding assays that allow the selection of aptamers was demonstrated^11, 26, 27^. However, this platform requires hardware modifications and additional software. Here, we present an approach that allows for performing HiTS-FLIP experiments with up to more than 100 binding assay’s cycles automated on a common sequencer without necessitating any hardware modifications or additional software. Using this innovative technology, we successfully selected aptamers for three proteins of clinical relevance for infections with *P. aeruginosa* (LecA, LecB, and PEA)^25, 28, 29^. The direct comparison with SELEX shows that our novel platform enabled the selection of higher affinity aptamers for each target with a significantly higher time-efficiency (one week instead of multiple months). The presented method also offers a high reproducibility with a mean coefficient of variation of 9.25% (mean standard deviation of 12 fluorescence units). The ease of implementing our method onto a MiSeq and its complete integrity with the regular sequencing procedure offers enormous potential to be widely implemented as an add-on to the already established NGS of aptamer pools^30^.

The binding affinities for selected aptamers determined by HiTS-FLIP showed overall high agreement with the results of established binding assays, including SPR and affinity chromatography. In addition, we validated that the fluorescence intensity or proximity to a FM cluster did not significantly influence the obtained HiTS-FLIP results. For example, the cluster PEA33764 had close vicinity to a FM cluster (Extended Data Fig. 5) and a low fluorescence intensity (*F*_max_ = 86). However, PEA37764 showed a similarly high affinity in the other performed assays (Table 1), implying that this did not influence the accuracy of the fluorescence intensity extraction by the MiSeq’s RTA, unlike earlier observations^11^.

In addition to selecting DNA aptamers for proteins and small molecules, our experimental setup also bears a great potential for studying the biophysics of naturally occurring DNA structures, such as G quadruplexes^31^ and the intercalated-motif^32^, and their interactions with different ligands, by displaying either synthetic oligonucleotides or genomic libraries.

Furthermore, HiTS-FLIP variations that enable studying binding to RNA or proteins with high-throughput on NGS flow cells have already been described. The procedures of those experiments (e. g. HiTS-RAP^33^, RNA-MaP^13^, Prot-MaP^34^, and deep screening^22^) differ from HiTS-FLIP by additional *in situ* transcription and translation following DNA sequencing. With this technology, these experiments can be performed in an automated manner on a MiSeq.

## Online Methods

### General Material and Methods

Other than the exceptions noted below, all chemicals and lab supplies were purchased from Sigma-Aldrich Corp. (St. Louis, MO, USA) or Carl Roth GmbH + Co. KG (Karlsruhe, Germany). The MiSeq sequencer as well as reagents required for sequencing (MiSeq Reagent Nano Kits v2 (300-cycle), Nextera XT DNA Library Preparation Kit, phiX Control v3) were purchased from Illumina Inc. (San Diego, CA, USA).

### Library design and construction

A PAGE-purified hand-mixed DNA library with corresponding primers as well as complementary primer-blocking oligonucleotides and fluorescently labelled FM-complements were purchased from Integrated DNA Technologies Inc. (Coralville, IA, USA). Primers were ordered with standard desalting while fluorescently labelled oligonucleotides where obtained HPLC-purified. A detailed listing of all oligonucleotides is provided in Table S1.

### Protein expression and purification

The gene sequences *lecA, lecB,* and *eta* of *P. aeruginosa* (coding for LecA (UniProt ID Q05097), LecB (UniProt ID Q9HYN5) and PEA (UniProt ID P11439), respectively) were optimised for the expression in *Escherichia coli* by the GeneOptimizer tool (GeneArt, Thermo Fisher Scientific Inc., Waltham, MA, USA) and manually revised using the codon usage database kazusa (www.kazusa.or.jp/codon/; an extended version of codon usage tabulated from GenBank, NCBI, last accessed on 01.05.2021). In order to generate a less toxic version of PEA, a mutant with R276G and E553D was encoded^35, 36^. The *lecA* gene was expressed by transformed *E. coli* ArcticExpress (DE3) cells while *lecB* and *eta* were expressed by transformed *E. coli* BL21 (DE3) cells (both strains from Agilent Technologies, Inc., Santa Clara, CA, USA).

All cells were electrocompetent and prepared according to the protocol of Dower et al.^37^. For transformation, between 10 pg and 100 ng of plasmid DNA (< 5 µL, pET-22b(+) vector encoding a C-terminal His_6_-tagged R276G and E553D PEA with an N-terminal pelB-leader, an N-terminal His_6_-tagged LecA, or an N-terminal His_6_-tagged LecB, all synthesised by GenScript Biotech Corp., Piscataway Township, NJ, USA) was placed in a sterile 1.5 mL cap and cooled on ice. In addition, electrocompetent cells were thawed on ice, 50 µL of them were transferred into the sterile pre-cooled 1.5 mL cap, homogenised by flicking and transferred as quickly as possible into an electroporation cuvette (gap width 0.2 cm) pre-cooled on ice. Transformation was performed by one pulse (25 µF, 200 Ω, 1.80 kV) using the Eporator (Eppendorf AG, Hamburg, Germany). Subsequently, the cells were immediately suspended in 1 mL of sterile SOC medium. This suspension was transferred to a sterile 2 mL cap and shaken for one hour at 37 °C and 200 rpm. The cells were afterwards spread on agar plates (LB agar with 100 mg/L ampicillin) and incubated at 37 °C overnight. Not transformed *E. coli* cells were used as a negative control.

For expression of the recombinant proteins, 100 mL of LB amp broth (100 µg/mL ampicillin) was inoculated with a single colony and shaken at 37 °C and 200 rpm in a shaking incubator until an optical density at 600 nm (OD_600_) of 0.4–0.6 was reached. This preculture was used to inoculate 800 mL of LB amp broth (100 µg/mL ampicillin) which was then incubated under shaking (150 rpm) until an OD_600_ of 0.4 was reached. Expression in *E. coli* BL21 (DE3) cells was induced by 0.5 mM IPTG at 22 °C and 130 rpm for 24 h while expression in ArcticExpress (DE3) cells was performed at 15 °C for 48 h. Cells were pelleted by centrifugation at 3,000 rcf and 3 °C for 30 min, washed once with 45 mL saline, centrifuged again at 3,000 rcf and 4 °C for 20 min, and stored at −80 °C.

An osmotic shock was performed according to a slightly modified protocol of Matthey et al.^38^ to disrupt only the periplasmic fraction of the cells, which contain the expressed PEA. For this purpose, the cell pellet was resuspended in 25 mL of ice-cold 5 mM MgSO_4_ by ultrasonication (< 30 s) and then incubated for 10 min on ice with periodical shaking. Subsequently, centrifugation was performed at 8,000 rcf for 20 min. The supernatant contained the periplasmic fraction.

To extract LecA and LecB, whole *E. coli* cells were disrupted. For this disruption, the cell pellets were resuspended in 3–5 mL of buffer A (5% glycerol, 15 mM imidazole, 0.04% (w/v, sodium azide, 500 mM sodium chloride, 50 mM Tris-HCl, pH 8.0; to which DNase 1 and 30 µM 4-(2-Aminoethyl)benzenesulfonylfluoride hydrochloride have been added) per gram of cells and afterwards dispersed by an ultrasonic homogeniser (< 30 s). Cell disruption was performed by a French press, pre-rinsed with buffer A, at 1.80 kbar under cooling to 4–8 °C and repeated once. The cell suspension was then centrifuged for 1 h at 4 °C with 27,000 rcf and the protein-containing supernatant was used for protein purification.

Exploiting the affinity of the fused His-tags, the recombinant proteins were purified by immobilised metal affinity chromatography. The corresponding protein-containing supernatant was separated at 8 °C using a nickel-chelate chromatography column within an ÄKTAPrime plus system (GE Healthcare, Little Chalfont, UK). Each buffer was membrane filtered (Nitrocellulose, 0.22 µm, 47 mm) and degassed before being applied to the column at a flow rate of 2 mL/min. For preparation of the column, the resin was resuspended in ddH_2_O, remaining Ni^2+^-ions were removed by 50 mL of 20 mM EDTA, followed by cleaning of the resin with 50 mL of 20 mM NaOH, and reactivation with 50 mL of 100 mM NiSO_4_. Afterwards, the column was rinsed with 160 mL of buffer A and the protein solution was applied to the column. The column was then rinsed again with 60 mL buffer A, and the expressed protein was eluted with an 80 mL-long stepless gradient from buffer A to buffer B (composition as buffer A but with 800 mM imidazole) and captured in 6 mL aliquots. The column was then rinsed with 50 mL buffer B followed by re-equilibration with 50 mL of buffer A.

The purity of the protein fractions was analysed by SDS-PAGE (8% gels for PEA, 16% gels for LecA and LecB) and Coomassie staining.

The protein fractions containing the largest amount and purest target protein were concentrated to a concentration of approximately 2 mg/mL using an Amicon stirred cell with a regenerated cellulose ultrafiltration membrane with a MWCO of 10 kDa. The concentrated protein solutions were then dialysed over night at 5 °C in 1 L of PBS and afterwards stored at −80 °C.

### SELEX

SELEX was performed slightly modified to the protocol described by Hünniger et al. (2014)^39^:

First, the target proteins were covalently coupled to SiMAG carboxyl magnetic particles with a diameter of 1.0 µm (50 mg/mL; chemicell GmbH, Berlin, Germany). For this, 10 mg SiMAG carboxyl magnetic particles were washed twice with 1 mL MES buffer (0.1 M, pH 5.0) and resuspended in 250 μL MES buffer with freshly added 10 mg EDC and agitated for 10 min at room temperature for activation. The activated beads were then washed twice with 1 mL MES buffer. Coupling was performed by adding 50–75 μg of purified protein to the activated beads and incubating for 2 h with gentle shaking at room temperature. After three more washes with 1 mL PBS each, the magnetic particles were blocked with 1 mL of blocking buffer. The target beads were washed with 100 μL double-distilled water (ddH_2_O) prior to use. The final concentration of particles was adjusted to 10 mg/mL, which corresponds according to the manufacturer’s instruction to 1.8 × 10^10^ beads/mL. Magnetic particles for the counter-SELEX were prepared in the same way, merely without target immobilisation to obtain just ethanolamine at the bead surface. Coupling of the His-tagged proteins was confirmed by an ELISA as described by Frohnmeyer^40^.

For the coupling of reverse primer sequences to magnetic beads, 25 µg of 5′-NH2−C12-reverse-primer (5′-/5AmMC12/-RP) was used. Primer coupling was carried out according to the protocol mentioned above but without the two MES washing steps after activation of the beads. In addition, the 50 μg of SELEX target were substituted with the primer and BSA (0.1%; w/v) was used as a blocking reagent in the blocking buffer.

The conditions applied for SELEX are summarised in Table S6–8. Before each target incubation, the corresponding aptamer pools were mixed with equimolar amounts of forward primer complementary oligonucleotides and reverse primers, then heated (5 min, 95 °C) followed by immediately cooling (0 °C, 10 min). The DNA pool was afterwards diluted with selection buffer. Incubation and washing steps during FISHing were performed automatically using the KingFisher Duo (Thermo Fisher Scientific Inc., Waltham, MA, USA) while the elution was performed manually. For the ‘heat’ elution, the beads with bound aptamers were suspended in 70 µL ddH_2_O, heated to 90 °C for 10 min and the eluate was separated using a magnetic rack. In the later selection rounds, the elution was performed competitively. For this elution, the beads with bound aptamers were suspended in 70 µL of ddH_2_O containing 1 mg/mL of the target protein, subsequently incubated for 60 min at 36 °C, and the eluate separated using a magnetic rack. To increase the stringency and thereby the efficiency, washing volumes were increased and incubation times were decreased during the SELEX process. The final elution step was performed in 70 µL ddH_2_O, so that the FISHing eluate could be used for the following BEAMing process without further preparation.

The amplification of the aptamer pool during each SELEX round was set up with a volume of 150 μL of aqueous phase in a 2 mL tube (15 µL of 10x DreamTaq buffer (Thermo Fisher Scientific Inc.), 9 µL of 0.5 U/µL DreamTaq polymerase (Thermo Fisher Scientific Inc.), 10 µL of 10 µM dNTPs, 4 µL of 100 µM FP, 2 µL of 5 µM RP, 6 µL of 5’-/5AmMC12/-RP beads, 30 µL of template DNA, 74 µL ddH_2_O). The aqueous solution was then overlaid with 46 µL ABIL WE 09 MB, 132 µL mineral oil, and 438 µL Tegosoft DEC (Evonik Industries AG, Essen, Germany) and subsequently homogenised by shaking for 5 min at the highest level at room temperature.

The reaction mixture was split into 80 μL aliquots and subjected to a PCR (initial denaturation for 5 min at 95 °C followed by 30 cycles of denaturation at 95 °C for 30 s, annealing at 64 °C for 30 s, and elongation at 72 °C for 30 s, followed by final elongation at 72 °C for 5 min and cooling at 4 °C). After PCR, the aliquots were combined, and the emulsion was broken by adding 1.5 mL of −21 °C isopropanol. The solution was then mixed until it was transparent. Subsequently, the reaction mixture was centrifuged for 5 min at 10,500 rcf to precipitate the magnetic beads and the supernatant was removed. The magnetic particles were resuspended in 80 µL ddH_2_O and the tube was incubated for 10 min at 95 °C for ssDNA elution. Immediately thereafter the still hot supernatant, which contained the desired aptamers, was removed in a magnetic rack and transferred to a new reaction tube. This supernatant represents the BEAMing eluate which is ready to be used for the next FISHing.

The yield of ssDNA was quantified after each BEAMing using a Quantus fluorometer (Promega Corp., Madison, WI, USA) in combination with the Quant-iT OliGreen ssDNA Assay Kit (Thermo Fisher Scientific Inc., Waltham, MA, USA) to set up DNA concentration for further SELEX rounds.

In addition, for the evaluation of each SELEX round, the quality of each BEAMing cycle was checked by a control-PCR (2.5 µL 10x DreamTaq buffer (Thermo Fisher Scientific Inc., Waltham, MA, USA), 0.5 µL 0.5 U/µL DreamTaq polymerase (Thermo Fisher Scientific Inc., Waltham, MA, USA), 2 µL of 10 µM dNTPs, 0.25 µL of 100 µM FP and RP each, 1 µL FISHing eluate, 18.5 µL ddH_2_O). 25 cycles of PCR were performed according to the temperature program given for BEAMing. A negative control (ultrapure water) and a positive control (aptamer or DNA library, 100 nM in ultrapure water) were always included. The control-PCR products were analysed by 3% agarose gel electrophoresis (TAE buffer, staining with GelRed, Genaxxon bioscience GmbH, Ulm, Germany) and UV visualisation. On observing DNA products with the expected length (90 bp), the BEAMing eluate was applied for the subsequent SELEX round.

Real-time PCR (qPCR) was used to determine to quantify the bound fraction of aptamers and to monitor their diversity decrease during SELEX. QPCR was performed in triplicate for selected selection rounds with the following Master mix: 2.5 µL 10x DreamTaq buffer, 0.5 µL of 0.5 U/µL DreamTaq polymerase (both from Thermo Fisher Scientific Inc., Waltham, MA, USA), 2 µL of 10 µM dNTPs, 0.15 µL each of 100 µM FP and RP, 1 µL of 1:1250 SYTO 9 Green (Thermo Fisher Scientific Inc., Waltham, MA, USA), 1 µL template, and 12.7 µL ddH_2_O. The qPCR and melting assay were performed by an initial denaturation at 95 °C for 5 min, 35 cycles of 95 °C for 30 s, 60 °C for 30 s, and 72 °C for 30 s, final elongation for 5 min at 72 °C and final denaturation at 95 °C for 2 min, re-hybridisation at 76 °C for 180 min and melting curve from 55 °C to 90 °C at a rate of 1 °C/15 s and subsequent cooling at 4 °C. A positive control (aptamer library, 1 nM) and a negative control (ddH_2_O) were always included. For absolute quantitation the aptamer library was analysed in triplicate in five different concentrations (100 fM, 1 pM, 10 pM, 100 pM, 1 nM).

### Library preparation for HiTS-FLIP

The aptamer pools for the HiTS-FLIP experiment were prepared according to the Illumina 16S metagenomic sequencing library preparation protocol^41^. Firstly, sequencing adapters (see Table S1) were attached to the chosen aptamer library by PCR. For this, 10 µL of Q5 Polymerase Buffer (5x, New England Biolabs Inc., Ipswich, MA, USA), 0.5 µL of Q5 High-Fidelity DNA Polymerase (New England Biolabs Inc., Ipswich, MA, USA), 1 µL of dNTPs (10 µM, biostep GmbH, Jahnsdorf, Germany), 5 µL of Fw and Rv Primer Seq adaptors (100 µM), 5 µL of template, as well as 23.5 µL of ddH_2_O were combined. A short random region was inserted between the Illumina sequencing primer and the aptamer FP to ensure enough base diversity in the first five cycles of sequencing for deconvolution of the different clusters of oligonucleotides. Furthermore, an EcoRI restriction site was included between the aptamers RP and the Illumina sequencing primer. This restriction site enables cleaving of the constant regions prior to interaction profiling to facilitate independent folding of the random region. The PCR was performed by 3 min of initial denaturation at 95 °C, followed by 4–12 cycles of 30 s each at 95 °C, 55 °C, and 72 °C, concluded by a final elongation for 5 min at 72 °C and subsequent cooling at 4 °C. As template for HiTS-FLIP experiments, pre-enriched aptamer pools (i. e. after three or five SELEX rounds) were used. A pilot PCR was performed to determine the correct number of cycles for amplification. For this, 10 µL of the reaction was taken after cycles 4, 6, 8, 10, and 12 and the amplicon was separated on a 3% agarose gel at 120 V for 45 min and visualised by staining with GelRed (Genaxxon bioscience GmbH, Ulm, Germany). For the final PCR reaction, the cycle number that yielded a product of the correct length without producing unwanted products was chosen. The desired amplicons’ bands were cut out from the gel and eluted using the Monarch Gel Extraction Kit (New England Biolabs Inc., Ipswich, MA, USA). The concentration of the library after the first PCR was determined fluorescently using the Quantus fluorimeter with the QuantiFluor dsDNA System (Promega Corp., Madison, WI, USA) in three-fold determination. The library was then amplified to approximately 30 nM while attaching Nextera XT indices. The composition of these indices PCR reaction was as follows: 5 µL DreamTaq Polymerase Buffer (10x, Thermo Fisher Scientific Inc., Waltham, MA, USA), 0.5 µL DreamTaq DNA Polymerase (Thermo Fisher Scientific Inc., Waltham, MA, USA), 1 µL of dNTPs (10 µM, biostep GmbH, Jahnsdorf, Germany), 5 µL of Nextera XT Index 1 Primer (N7XX) and Nextera XT Index 2 Primer (S5XX), 5 µL of template, as well as 28.5 µL of ddH_2_O. The PCR conditions corresponded to those of the adapter PCR but with 21 cycles. The desired amplicons were again purified as described for the first PCR.

The correct length of the amplicons was analysed via Bioanalyzer using a High Sensitivity DNA kit (Agilent Technologies, Inc., Santa Clara, CA, USA) and quantified by fluorescence as described above.

The pooled libraries (for PEA’s HiTS-FLIP: 65% PEA3, 15% FM oligo, 0.005% R14.33, 20% phiX; for LecA’s and LecB’s HiTS-FLIP: 39.25% LecA5, 39.25% LecB5, 1.5% FM, 20% phiX) were denatured and diluted to a final concentration of 6 pM as given in Illumina’s protocol.

### Protein labelling with AF647

The purified target proteins were labelled with Alexa Fluor 647 (AF647) amine-reactive, activated NHS esters (Thermo Fisher Scientific Inc., Waltham, MA, USA and Lumiprobe GmbH, Hannover, Germany). This fluorophore aligns with the ‘A’ channel of the MiSeq-system.

For labelling with AF647, the purified protein was concentrated to > 2 mg/mL using Amicon centrifuge inserts with a MWCO of 10 kDa (Merck Millipore Corp., Billerica, MA, USA) and dialysed with 10 mM PBS. To adjust to an alkaline pH value, the solution containing 0.5 mg of the protein was mixed with 5% (v/v) of NaHCO_3_ buffer (1 M, pH 8.5). An AF647 solution was prepared at a concentration of 10 mg/mL in anhydrous DMSO. 5 µL of this dye solution was added to the protein solution under constant mixing and then incubated for 60 min with continuous mixing (600 rpm) at room temperature. Unbound fluorophore was afterwards removed by four-fold dialysis with PBS at 4 °C using the Pur-A-Lyzer Midi Dialysis Kit (MWCO 3.5 kDa, Merck Millipore Corp., Billerica, MA, USA).

The labelling efficiency was determined photometrically with the NanoDrop One^C^ (Thermo Fisher Scientific Inc., Waltham, MA, USA). A labelling efficiency in between 0.75 and 2 was considered appropriate for the HiTS-FLIP experiments.

### Hardware modification of the Illumina MiSeq

As the first option, the hardware of the MiSeq was minorly modified to perform the HiTS-FLIP experiment. In order to deliver additional reagents (i. e. fluorescently labelled target, formamide, pre-imaging buffer, and hybridisation buffer) to the flow cell, an external valve (C25Z-31812EUHB, Valco Instruments Co. Inc., Houston, TX, USA; VICI) was connected to port 23 of the sequencer’s internal valve (a C25G valve from VICI), as outlined in Fig. S1. Compatible tubing and fittings were purchased from Cole Parmer GmbH (part no. 06407-41 and F-130, Wertheim, Germany) as well as from VICI (part no. CNNF1PK). From port 23, the tubing used for the connection was routed through an existing hole in the bottom of the instrument and connected to the external valve in front of the MiSeq (Fig. S1B). Proprietarily, port 23 is operated solely to generate an air bubble to physically separate two solutions during washing of the instrument and during measurement of the flow but is not used during sequencing. To ensure that this functionality remains, the external valve is automatically switched to an unoccupied position (i. e. 1) every time the system is started. Furthermore, ports 18–20 are proprietarily designated to custom primers and thus not necessarily used in the actual sequencing flow. Hence, they are also available for the HiTS-FLIP experiment. To the ports 18–20 reagents are supplied via the regular cartridge. An exemplary assignment of the external valve as well as ports 18–20, as applied during the PEA-HiTS-FLIP experiment is listed in Table S2.

### Software modification of the Illumina MiSeq for application with additional external valve

The software of the MiSeq was modified to perform the fluorescent ligand interaction profiling during the ‘second read’. For this, the folder structure of the MiSeq marked in dark blue in the organisation chart (Fig. S2) was newly created. The function and contents of the newly created folders are described in the GitHub project. A new recipe (Fig. S2, ‘HiTS_FLIP_Recipe’) was created by adding ‘ArrayGen’, ‘CompleteCycleFLIP’, ‘EndReadFLIP’ to ‘Chemistry’ and modifying ‘Reads’ as well as ‘Sequencing’ to change the procedure so that no sequencing is performed in the second read, but instead the target protein is applied to the flow cell at different concentrations. Additionally, the integration times in ‘Sony’ were adapted to the needs of the HiTS-FLIP experiment. Furthermore, the CFG files ‘MiSeqOverride’, ‘MiSeqSoftware.Options’ and ‘MiSeqCommon’ were modified in order to choose the focus parameters according to the fluorophores used for the FMs as well as for the labelled proteins. The detailed modifications and xml files are provided upon request. Python programming language version 3.8.5 (Python Software Foundation, https://www.python.org/) was installed to control the external valve. To facilitate the installation of Python packages, the package management program pip (the pip developers, https://pypi.org/project/pip/) has been installed. The installed packages and their usage are listed in alphabetical order in Table S5. The communication with the valve was performed by means of the Python module ViciValve (Sebastian Steiner, https://pypi.org/project/vicivalve/, version 0.0.12).

Our automated valve control takes advantage of the MiSeq’s ability to store all actions performed in log files in real-time, such as optics settings, valve movements, or pump commands. These files can be read at regular intervals (e. g. of approx.1 s) to track which task the MiSeq is currently performing, and it is possible to trigger an action in response to certain log entries. For this approach we utilised the function WaitCommand() used in the chemistry xml files to be able to set reaction times. Since the time is specified in milliseconds and Illumina solely uses whole seconds, the last three digits are proprietarily always 000. If the device waits, the waiting time is also recorded in the log files and thus the insertion of a specific time in the xml files represents a possibility to, for example, switch the valve at a specific time in the course of the run. At the same time, the duration can be adjusted to allow enough time to recognise the entry in the log file, decode the number into a position, and switch the valve.

The times we used for this process start with the digits ‘135’, in order to differ as much as possible from the response times applied by Illumina in the code. WaitCommands 13501–13512 encode switching the external valve to the position of the last two digits while 13520 switches the external valve to a cycle dependent position specified in an ini file (Fig. S2, ‘HiTS_FLIP_INI). These commands are recognised by the script ‘HitsFlipMain.py’ (Fig. S2, ‘PythonCode’) which is provided upon request.

### Software modification of the Illumina MiSeq for application without hardware modifications

To perform a HiTS-FLIP without hardware modifications, the experiment was split into two parts. At first, regular sequencing of the library with a subsequent paired-end turnaround was performed. Secondly, the FLIP was executed in a second run on the MiSeq. For this purpose, a used reagent cartridge was disassembled, rinsed, and filled with custom reagents (Table S3). To prevent the consumables from being reused, Illumina equipped all sequencing consumables with a RFID-chip, which is timestamped and locked at the beginning of cycle 2 of the sequencing. However, if cluster generation fails during a run, the reagents are consumed but the RFIDs are still blank. Otherwise, Illumina can be contacted in order to receive RFID bypass codes. For the presented HiTS-FLIP experiment with LecA and LecB, we utilised the three consumables of a failed previous run to unlock the MiSeqControlSoftware during setup and exchanged them for the custom-filled cartridge, as well as for the flow cell and the PR2 bottle used during sequencing. A completely new recipe was designed to utilise the first read starting with a FirstBase for performing an autofocus followed by enzymatic digestion and triplicate incubation of 10 different protein concentrations and 10 blanks. The required xml files are provided upon request.

### Sequencing of the DNA libraries

The sequencing of the DNA libraries was performed using MiSeq Reagent Nano Kits v2 (300-cycles) with 105 cycles in read 1. During sequencing, the HiTS_FLIP_Main.py script was started, and the external valve primed manually, if not done earlier.

### Generation of the aptamer array

For the generation of the aptamer array after sequencing and the paired end turnaround, first, EcoRI digestion was used to remove the 3’ constant regions (adaptor, index, and flow cell primer sequences) in order to prevent potential steric hindrance between the aptamer and the target and to improve independent folding of the aptamers. By hybridising a complementary strand to the EcoRI recognition sequence (Rv Primer Seq Adapter) a double-stranded cut-site was formed that enables enzymatic restriction.

EcoRI-HF and rCutSmartBuffer were purchased from New England Biolabs Inc. (Ipswich, MA, USA). To validate the successful restriction, we incubated a flow cell with fluorescently labelled EcoRI cy5 Rv adapter (Table S1), which is only able to bind to the clusters if restriction was not successful (Extended Data, Fig. 4).

Furthermore, we hybridised the constant region of the aptamers with complementary oligonucleotides (FPc and RP, Table S1) to enhance independent folding of the random region.

### Protein binding experiments on the aptamer array

To carry out the HiTS-FLIP experiment using the external valve, the device was placed in front of the optical bench of the MiSeq. The external reagents were stored on cold packs (–18 °C) during the experiment (Fig. S1A). Before the interaction profiling could be started, the external valve was primed to ensure that all tubes are filled with reagents up to the valve without air bubbles. This step can be performed manually even during the sequencing process.

For control of the external valve, HitsFlipMain.py had to be started after beginning of the sequencing. The execution of the script starts immediately without any further input, so the ini file and the sample sheet must have been updated in the respective folders before starting.

If the HiTS-FLIP was performed using the custom-filled cartridge, the script can be started straightforwardly, as the recipe already includes a priming procedure.

### Alignment, pre-processing of images, and fitting of binding curves

After sequencing, the MiSeq saves location of the clusters along with the sequence and Q-score, the position and the intensities in fastq, locs, and cif files respectively. The intensities and the sequencing data were conjoined and saved in one csv file per channel using a script of Wu et al.^11^, which we modified to also deliver the Q-Scores for further quality assessment. The modified script will be provided upon request alongside with an application example for the HiTS-FLIP for LecA and LecB.

For quality filtering and curve fitting MATLAB Version 2023a (The MathWorks Inc., Natick, MA, USA) was used to first remove all intensities over 6000 A.U., as these are commonly induced by saturated pixels^13^. The sequences were deprived of their primer sequences and aptamers two bases shorter or longer than the N_50_-region of the initial library and sequences with more than three bases with a Q-score < 30 were omitted. Subsequently a HILL-function as given in Equation 1 was fitted using MATLABs fitnlm-function. A HILL-coefficient of 1 was applied as the majority of aptamers most probably exhibit one binding site only.

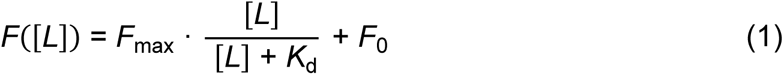

To evaluate the fits’ quality, the root mean square error (RMSE) was normalised to *F*_max_, resulting in the normalised root mean square error (NRMSE). Fits with an NRMSE exceeding 0.1 or a *K*_d_ falling outside the range of 0.1 nM to 2000 nM were excluded. Since most sequences were displayed only once, further statistical analysis was deemed unnecessary. Instead, clusters were sorted based on *K*_d_ and visually examined to identify any anomalies. To eliminate potential artifacts, such as interference with FM or phiX-clusters, the raw images were aligned with the sequencing data. Clusters were then selected based on their position in the FASTQ-files and subjected to visual inspection. The used code will be provided upon request.

### DNA secondary structure predictions

Initially, the secondary structure of the aptamers was forecasted using mfold^42^ under experimental conditions. Following this, RNAComposer server^43^ was employed to predict the most stable tertiary structure based on the inferred secondary structure. Next, HDOCK server^44^ facilitated the *ab initio* molecular docking of the aptamers’ 3D structure with their target proteins.

### Determination of dissociation constants by surface plasmon resonance spectroscopy

All SPR assays were conducted utilising the Sierra SPR-32 Pro system and biotin-tag capture sensors (BTC) from Bruker Daltonics GmbH & Co. KG (Bremen, Germany) at 25 °C. In our aptamer assay, two of the four spots per channel were allocated for aptamer interaction analysis, while the remaining two served as negative controls for blanking, displaying the chip’s neutravidin surface or the single stranded capture oligonucleotide (i. e. biotinylated RP).

For preparation of the sensor in the reversible aptamer assay, all spots of a BTC SPR sensor underwent preconditioning through three cycles with injections of 0.1 M HCl followed by PBS with 0.05% Tween20 (pH 7.4, PBST) at a flow rate of 25 μL/min, with each injection comprising a 30 s contact time and a 10 s dissociation time. Subsequently, the sensor was washed with PBST at 15 μL/min and a 1 min contact time. Following these washes, 50 nM of 5’ biotinylated RP for the aptamer library (Table S1) was permanently captured on the BTC sensor by incubation at 10 μL/min and a 5 min contact time, addressing spot B first, then C, and finally D, to ensure uniform capture levels. To eliminate loosely bound oligonucleotides, the three HCl washing steps were repeated. Finally, all spots were washed with PBST at 15 μL/min and a 2 min contact time.

One cycle of the subsequent binding analysis commenced with surface regeneration using 0.1 M HCl, followed by equilibration using running buffer (PBST + 1 mM MgCl_2_) at 25 μL/min and a 1 min contact time each. First, only spot B of each channel was addressed, and the aptamer was applied at concentrations of 10–100 nM with a flow rate of 10 μL/min for 3 min. This process was then repeated for the spot D under the same conditions. Subsequently, two analyte injections were carried out at a flow rate of 40 μL/min, with a 4–5 min contact time and a 5 min dissociation time, addressing all spots: first, the running buffer as a blank, followed by the target protein at various concentrations. Each cycle concluded with surface regeneration using 0.1 M HCl at 25 μL/min and a 1 min contact time. This procedure was repeated until all protein concentrations were analysed. Through the presence of ssDNA primers on spot C, the assay inherently integrates a negative binding control, evaluating nonspecific interactions between the target molecule and ssDNA.

Data analysis was performed using Bruker’s ‘SPR Analyzer 4’ software, involving a double subtraction utilising both the buffer injection and the signal of reference spot C of the same injection cycle. Additionally, a median filter was applied, followed by global kinetic fitting using a 1:1 Langmuir model.

### Native nano electrospray ionisation mass spectrometry

For nano-ESI-nMS, the protein was dialysed four-fold in 150 mM ammonium acetate. In those cases where the protein was to be analysed in complex with an aptamer, a concentration of 12.5 μM protein was incubated with 25 μM aptamer for a minimum of 30 min at room temperature before injection into the nano-ESI source.

Sample aliquots (2 µL) were loaded into custom-made nano-ESI gold-coated capillaries, affixed to the nano-ESI source, using a microliter syringe attached to flexible fused silica tubing. The measurements were conducted using a nano-ESI quadrupole time-of-flight instrument (Q-TOF2, Micromass/Waters, MS Vision, Waters Corporation, Milford, MA, USA) modified for higher masses. The parameters for the nano-ESI-nMS measurements can be found in Table S9. Samples were ionised in positive ion mode, with the pressure in the source region maintained at 10 mbar throughout. For desolvation and dissociation, the pressure in the collision cell was adjusted to 1.3–1.5 × 10^−2^ mbar argon In the ESI-MS overview spectra, the quadrupole profile spanned from 100 to 10,000 m/z.

### Determination of dissociation constants by fluorescence assay after bead-based affinity chromatography

The *K*_D_ values were not only determined by HiTS-FLIP and SPR, but also using affinity chromatography as described previously by Fischer et al.^45^. The target protein was immobilised on the surface of carboxylated magnetic particles (chemicell GmbH, Berlin, Germany) as described in the Section ‘SELEX’. Fluorescent 5′ 6-FAM-labelled oligonucleotides were diluted in a total volume of 90 µL with PBS to, 25 nM, 50 nM, 150 nM, 300 nM, 450 nM, 900 nM, 1.35 µM, 4 µM, and 12 µM. Afterwards, 10 µL of target beads (10 mg/mL) were added to each dilution and the resulting solution was incubated for 60 min at room temperature under mild shaking and light exclusion. The supernatant was removed, the beads resuspended in 100 μL ddH_2_O and transferred into a microtiter plate for fluorescence measurement (SpectraMax2; extinction 495 nm, emission: 520 nm, Molecular Devices, LLC. San Jose, CA, USA). Every experiment was performed in triplicate and blank measurements were included. Data were fitted non-linear (HILL-fit) using OriginPro 2019 software (OriginLab Corp., Northampton, MA, USA).

### Immunohistochemistry of LecB with human nasal concha tissue

LecB was biotinylated after a two-fold dialysis into amine-free buffer (PBS) by adding 100 μg biotin–NHS ester (10 mg/mL in water-free dimethyl formamide) to 1 mg of protein (1 mg/mL) in a glass vial. Subsequently, the reaction mixture was incubated under constant mixing and shaking at 400 rpm for 2 h at room temperature.

Biotinylation was confirmed via dot blot, for which 10 μL of the biotinylated protein was spotted onto a nitrocellulose membrane (pore size 0.45 μm). After drying, the membrane was blocked with 20 mL of TBS containing 0.2 g of BSA for 30 min. Subsequently, the membrane was incubated for 30 min with 20 μL of streptavidin-HRP in 10 mL TBS, followed by staining with the HRP substrate 3,3’-diaminobenzidine until desired development was achieved (∼10 min).

Sections of human nasal concha tissue were prepared through paraffin sectioning, followed by deparaffinization and pre-conditioning with TBS-based lectin buffer over three rounds of 5 min incubations.

For lectin immunohistochemistry, the biotinylated lectin was heated to 37 °C in TBS-based lectin buffer, then treated with 0.1% trypsin and incubated for 10 min at 37 °C. The biotinylated lectin was then diluted to 10 μg/mL in TBS-based lectin buffer and incubated with tissue sections for 30 min at room temperature. For inhibition analysis, lectins were incubated with different concentrations of monosaccharides or aptamers for 30 min prior to tissue incubation. TBS-based lectin buffer without proteins served as a negative control, always analysed concurrently. Following lectin-tissue section incubation, sections were washed twice for 5 min with TBST and once for 5 min with TBS. Bound lectins were detected using the Vectastain ABC Kit AK5000 (Vector Laboratories Inc. Newark, CA, USA) and Zytomed Permanent AP-Red Kit (Zytomed Systems GmbH, Berlin, Germany) following the manufacturers’ instructions. Sections were stained by adding Permanent Red and incubating for about 20 min at room temperature, followed by counter-staining with hematoxylin for 2–8 s at room temperature. Finally, sections were dehydrated using an increasingly concentrated ethanol series and permanently covered.

## Supplementary Information

Figure S1: Hardware modifications of the MiSeq; Figure S2: Characterisation of the recombinant PEA protein; Figure S3: Characterisation of the recombinant LecA protein; Figure S4: Characterisation of the recombinant LecB protein; Figure S5: Assessment of performed SELEX processes; Figure S6: HiTS-FLIP for PEA; Figure S7: HiTS-FLIP for LecA and LecB; Figure S8: Sensorgrams of the interaction of PEA aptamers; Figure S9: Affinity chromatography for PEA aptamers; Figure S10: Nano-ESI-nMS of PEA aptamers; Figure S11: Affinity chromatography for LecA and LecB aptamers; Figure S12: Predicted secondary structures of all aptamers

Table S1: Sequences of oligonucleotides used; Table S2: Assignment of external valve; Table S3: Assignment of custom-filled cartridge; Table S4: Newly created folders of the MiSeq; Table S5: Installed Python packages; Table S6: FISHing parameters of LecA-SELEX; Table S7: FISHing parameters of LecB-SELEX; Table S8: FISHing parameters of PEA-SELEX; Table S9: Nano-ESI-nMS parameters.

## Credit author statement

Conceptualisation, Resources, Supervision, Funding acquisition: AD, CU, ZI, US, MF; Project administration: AD; Methodology, Software, Validation, Data Curation, Writing – Original Draft, Visualisation: AD, CA; Investigation: AD, CA, TK, NE, AFRG, US; Writing – Review & Editing: TK, NE, AFRG, CU, ZI, US, MF

## Data Availability Statement

All extracted fluorescence data already linked to the clusters’ sequence is provided in the Open Science Framework repository under the following link: https://osf.io/mqsyd/?view_only=ca5955427b954a02be4a6782e574153f.

## Acknowledgements

The authors want to thank Lukas Jarren, Stephan Seifert, Nils Wax, and Maike Arndt for providing expertise.

Special gratitude is extended to to Martin Schwinzer for his support and expertise with the CD measurements, and to Maike Märker for performing the lectin immunohistochemistry experiments. Furthermore, we want to thank the SPR team of Bruker Daltonics, in particular Sven Malik and Paul Ritter, for providing us access to their SPR-32 Pro as well as for their support during the aptamer assay’s development. Nasal concha tissue was provided by Prof. Dr. med. Andreas Gocht (University Hospital Schleswig-Holstein, Germany).

## Extended Data

**Figure 4:**
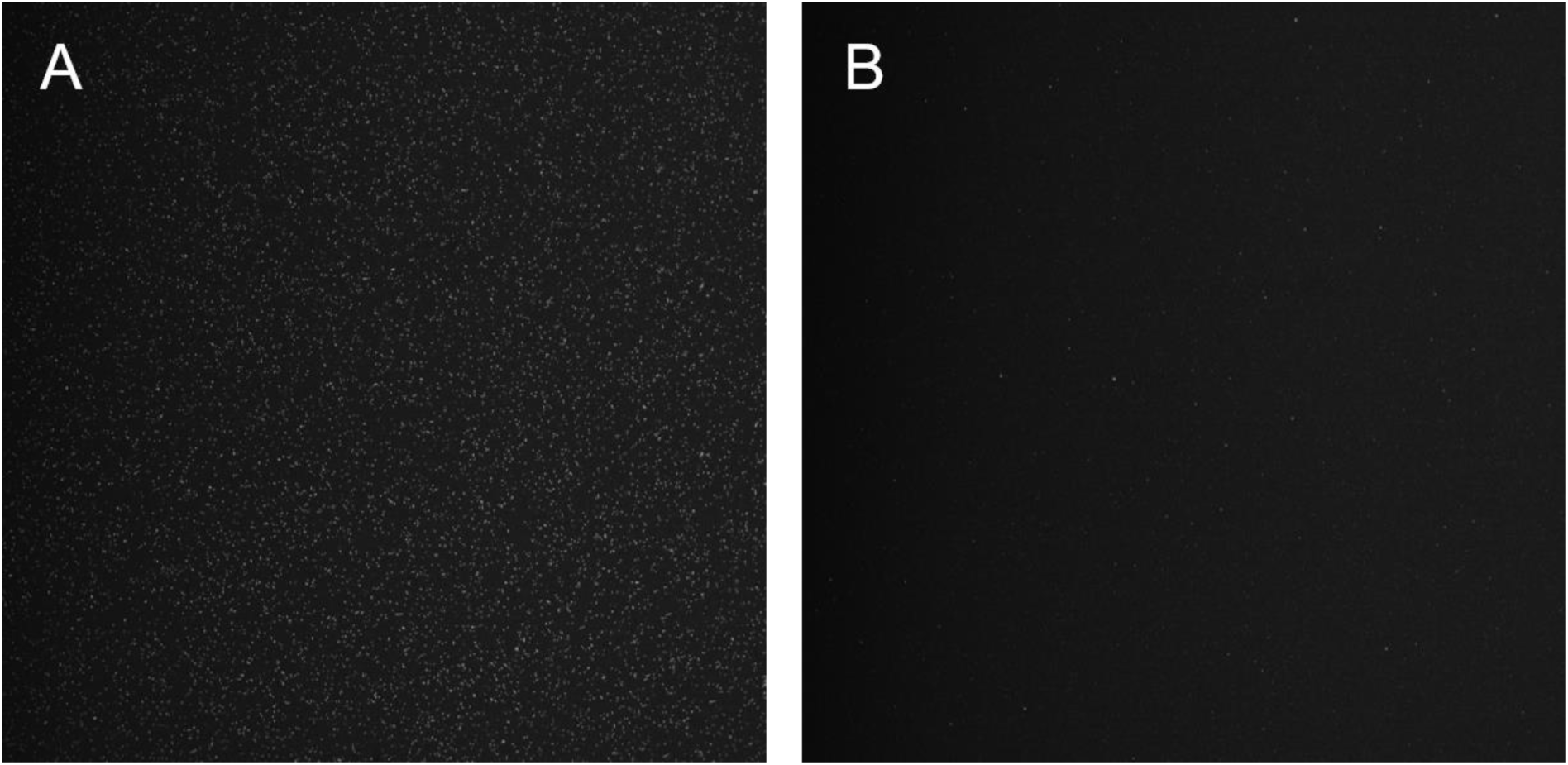
Images of a flow cell incubated with EcoRI cy5 Rv adapter oligonucleotide before (**A**) and after restriction with EcoRI (**B**). The images were acquired using the ‘A’ channel of the MiSeq and contrast adjusted by the same factor.

**Figure 5:**
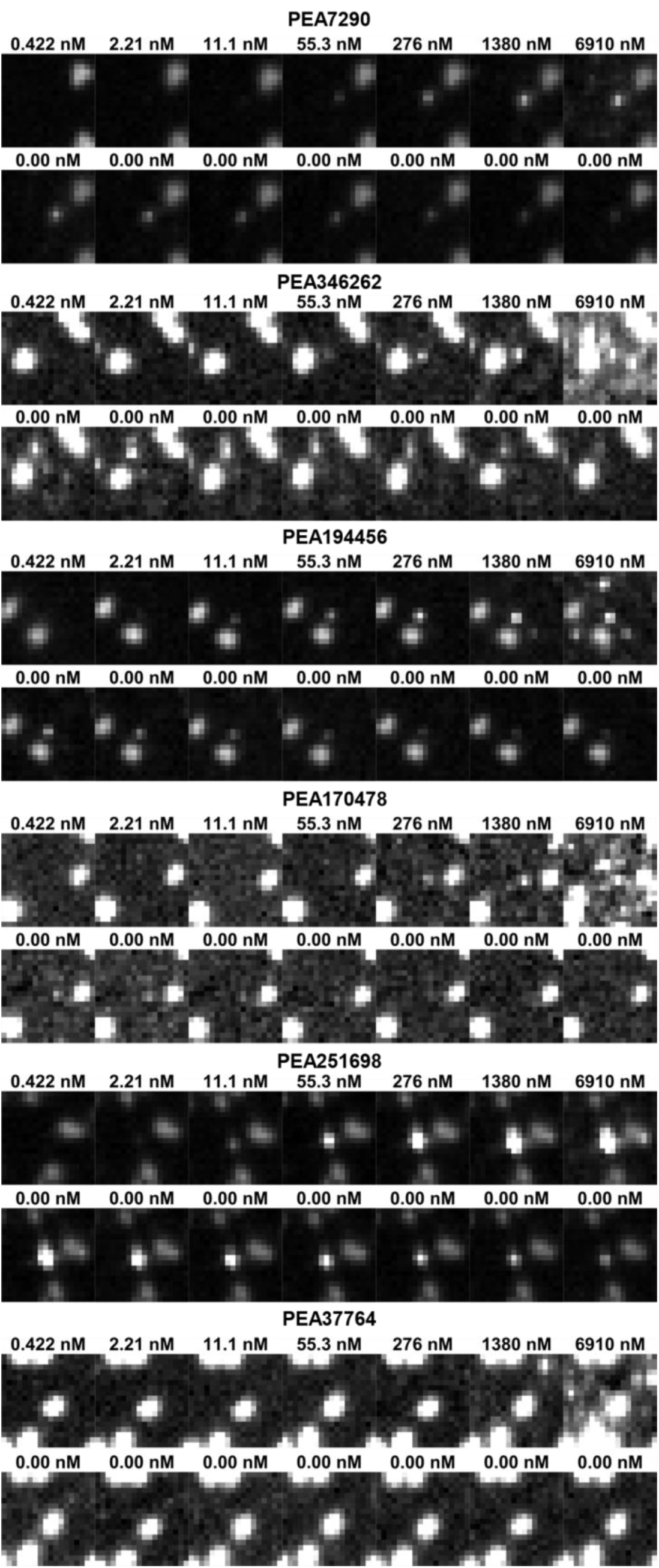
Images of selected clusters of PEA aptamers displayed on the flow cell during the binding assay.

