## Supplementary Figures and Tables for "Automated high-throughput selection of DNA aptamers using a common optical next-generation sequencer"

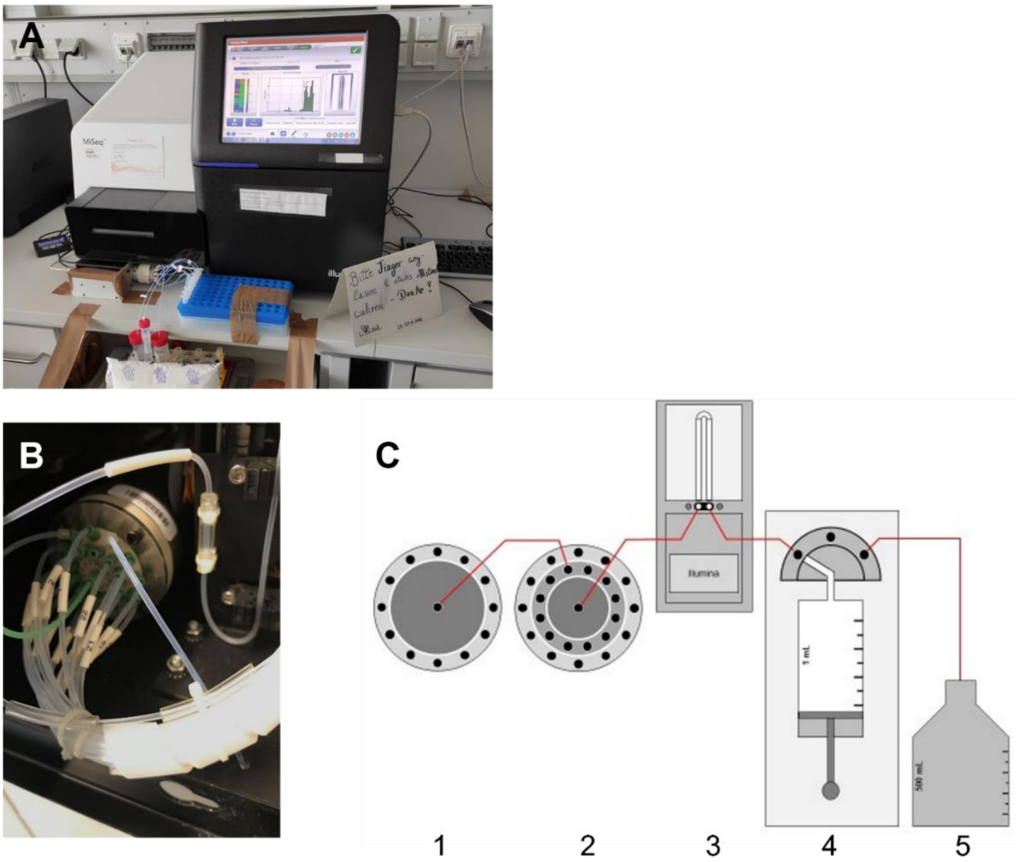


**Figure S1:** Hardware modification of the Illumina MiSeq. (**A**) Picture of the modified MiSeq with the external valve placed in front of the sequencer during a HiTS-FLIP experiment. (**B**) The new connection of port 23 of the internal valve to the external valve. (**C**) Schematic of the modified fluidics layout of the MiSeq. (1): External valve (VICI), (2): internal valve (VICI), (3): flow cell (Illumina), (4): syringe pump (Tecan Cavro), (5): waste container. For clarity, the connections to the reagent cartridge are not shown.

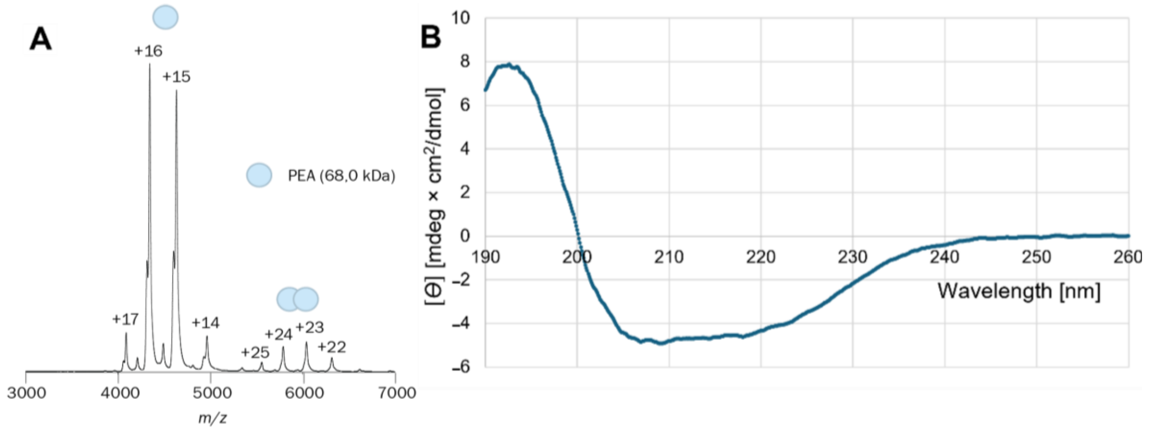


**Figure S2**: (**A**) Native mass spectrum of the recombinant PEA protein (MW = 68.0 kDa) confirming the predominant monomeric structure (*m/z* 4000–5000) and low abundance of dimers (*m/z* 5500–6500). (**B**) CD spectrum of the recombinant PEA protein recorded at a protein concentration of 0.09 mg/mL.

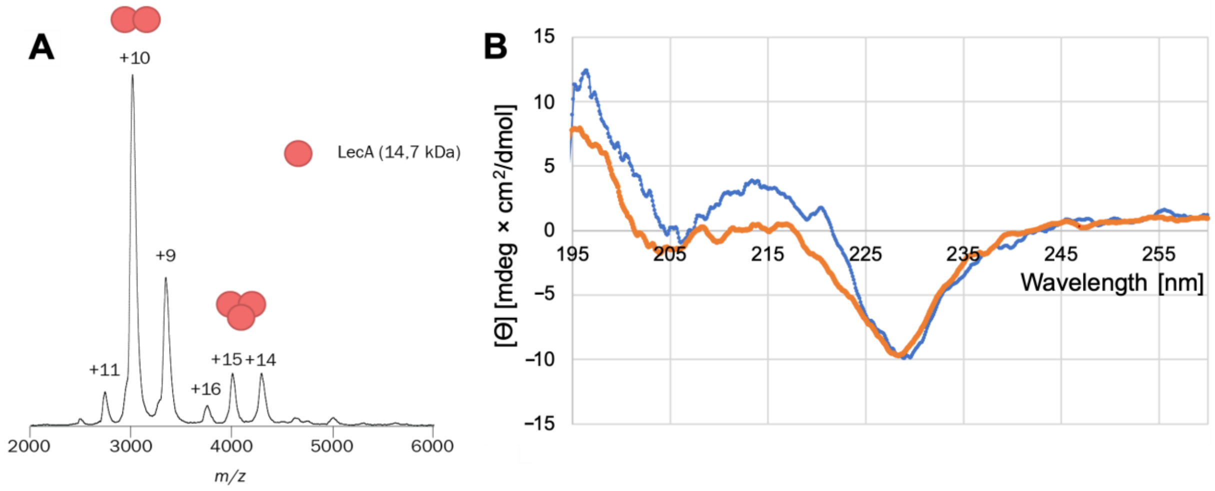


**Figure S3**: (**A**) Native mass spectrum of the recombinant LecA protein (MW = 14.7 kDa) confirming the formation of dimers. (**B**) CD spectra of the recombinant LecA protein (blue) and of the reference protein (orange) recorded at a protein concentration of ~0.1 mg/mL.

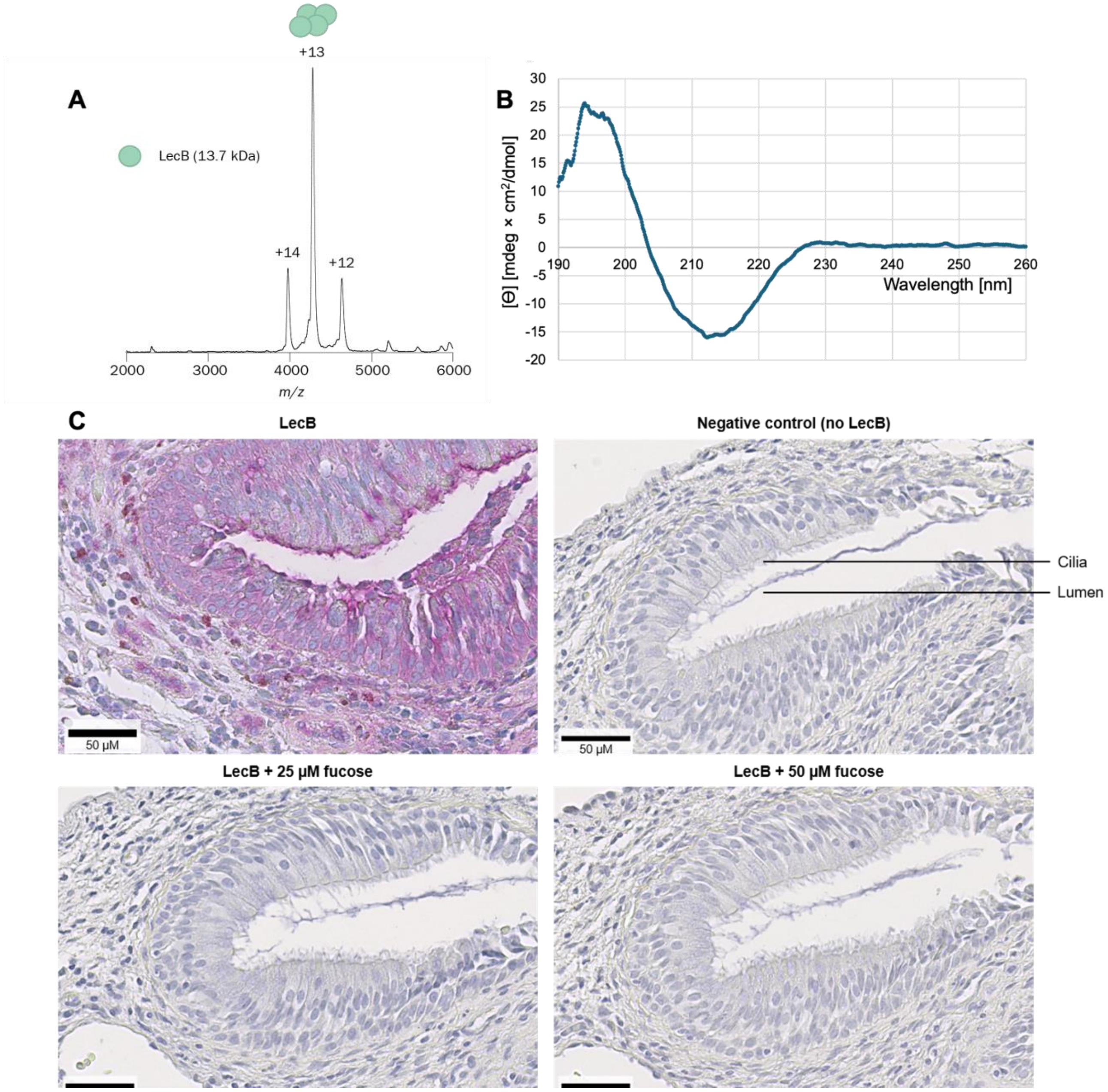


**Figure S4**: (**A**) Native mass spectrum of the recombinant LecB protein (MW = 13.7 kDa) confirming the formation of tetramers. (**B**) CD spectrum of the recombinant LecB recorded at a protein concentration of 0.11 mg/mL. (**C**) Immunohistochemistry showing the binding of recombinant LecB protein to human nasal concha tissue. Stained sections are shown after incubation without LecB (negative control), with the protein alone, as well as with a complex of LecB together with 25 µM or 50 µM L-fucose.

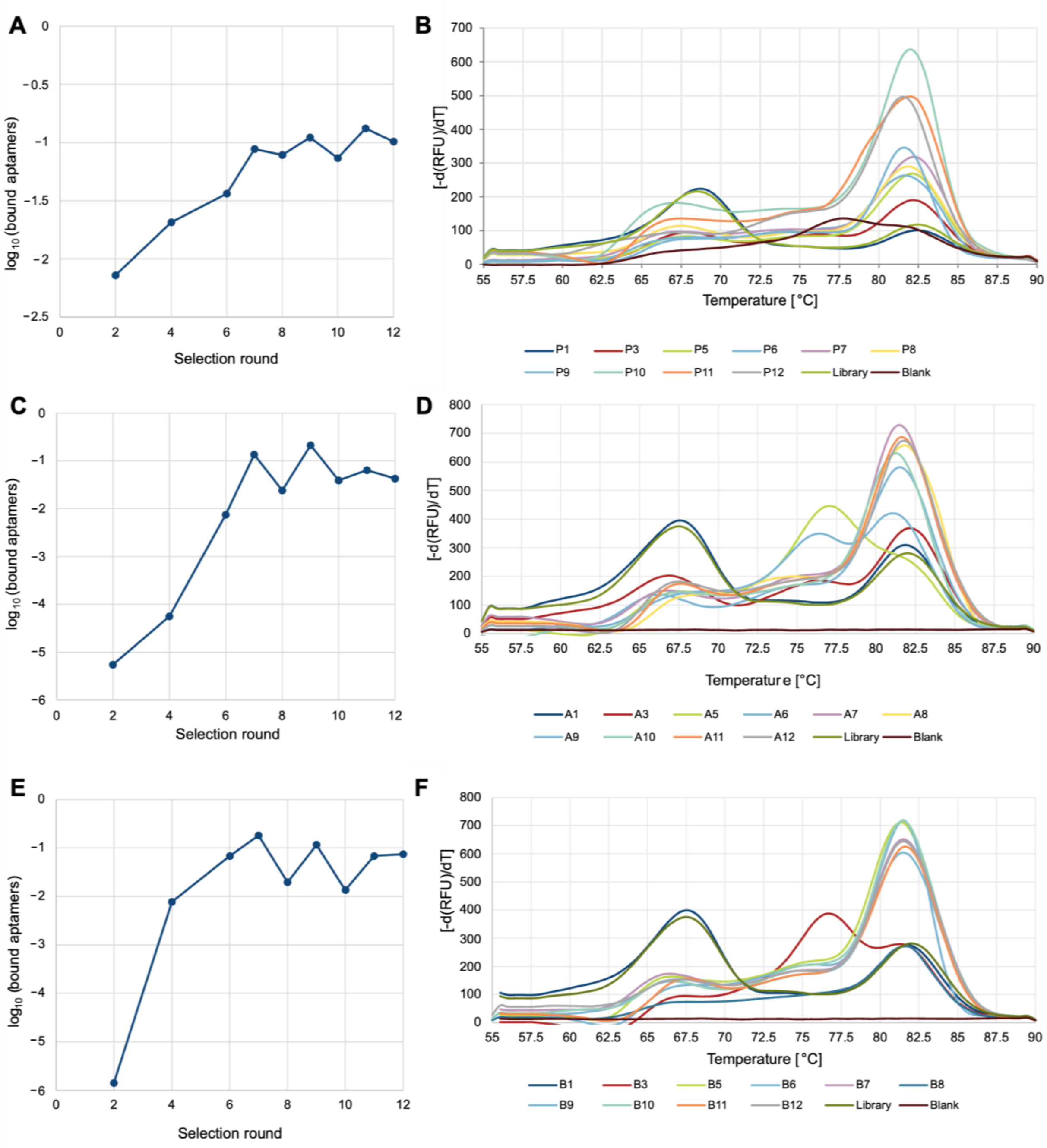


**Figure S5**: Assessment of target-bound aptamer percentage in successive rounds of the SELEX' for PEA (**A**), LecA (**C**), and LecB (**E**) using the calculated $C_t$ values obtained by qPCR.Melt curve analysis for the assessment of the SELEX processes for PEA (**B**), LecA (**D**), and LecB (**F**) in qPCR. P1–P12: BEAMing eluate of the respective PEA-SELEX rounds. A1–A12: BEAMing eluate of the respective LecA-SELEX rounds. B1–B12: BEAMing eluate of the respective LecB-SELEX rounds.

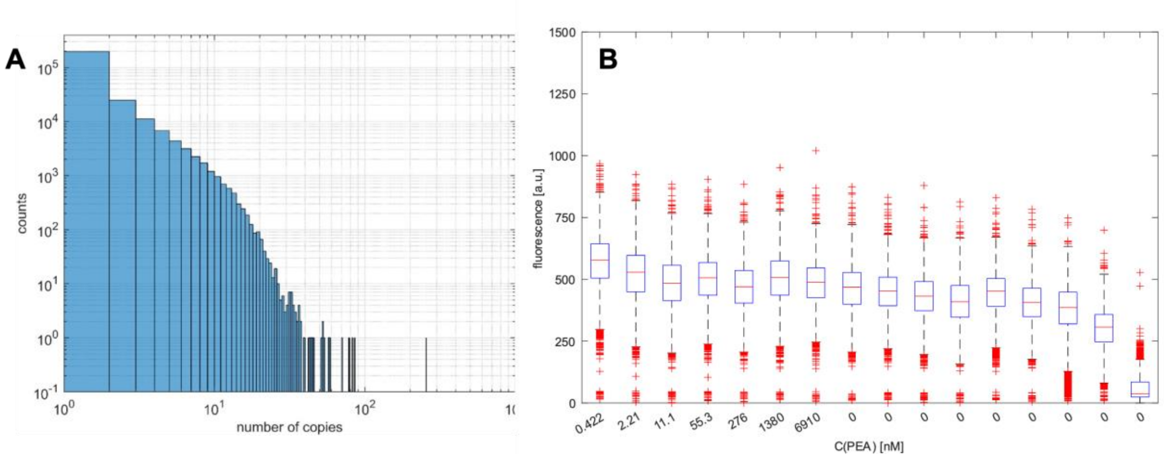


**Figure S6**: (A) Redundancy of sequences within the clusters displayed during HiTS-FLIP for PEA using a library pre-enriched by three SELEX cycles. (B) Box-whiskers plot of the fluorescence of FM-A throughout the binding assay in PBST + 1 mM $MgCl_2$ with an illumination time of 1200 ms. Outliers were defined as a value more than 1.5 times the interquartile range away from the bottom or top of the box.

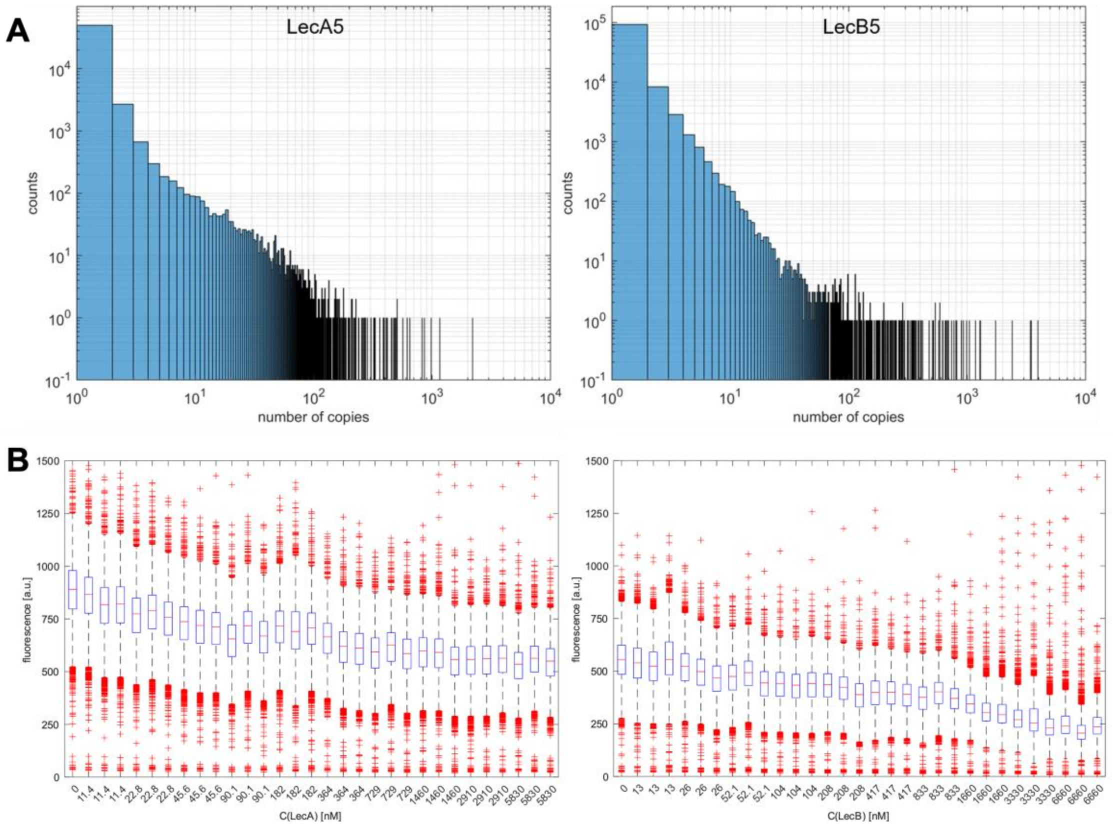


**Figure S7**: (**A**) Redundancy of sequences within the clusters displayed during HiTS-FLIP for LecA and LecB using a combined library of two pools pre-enriched by five SELEX cycles for one of the targets each. (**B**) Box-whiskers plot of the fluorescence of FM-A throughout the LecB and LecA binding assays in Tris-based lectin buffer with 0.05% Tween20 with an illumination time of 2500 ms. Outliers were defined as a value more than 1.5 times the interquartile range away from the bottom or top of the box.

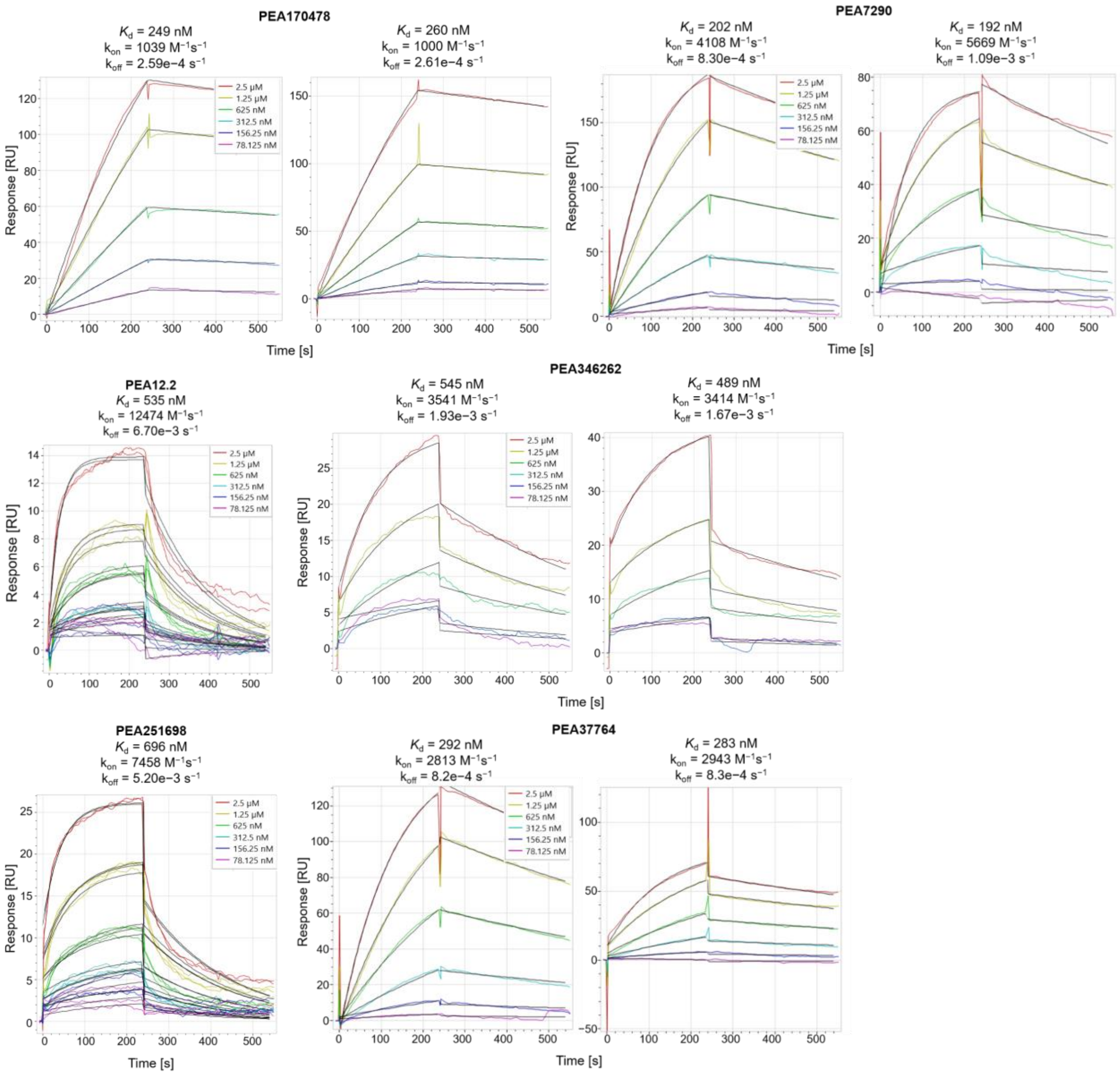


**Figure S8**: Sensorgrams of the interactions of PEA aptamers with PEA. The given sensorgrams have been recorded at aptamer capture concentrations of 25 nM (for PEA251698, PEA346262, PEA12.2), 50 nM (for PEA7290 right and PEA37764 right), or 100 nM (for PEA170478, PEA7290 left, PEA37764 left). All sensorgrams have been subjected to double subtraction using the blank of the same cycle as well as reference spot C.

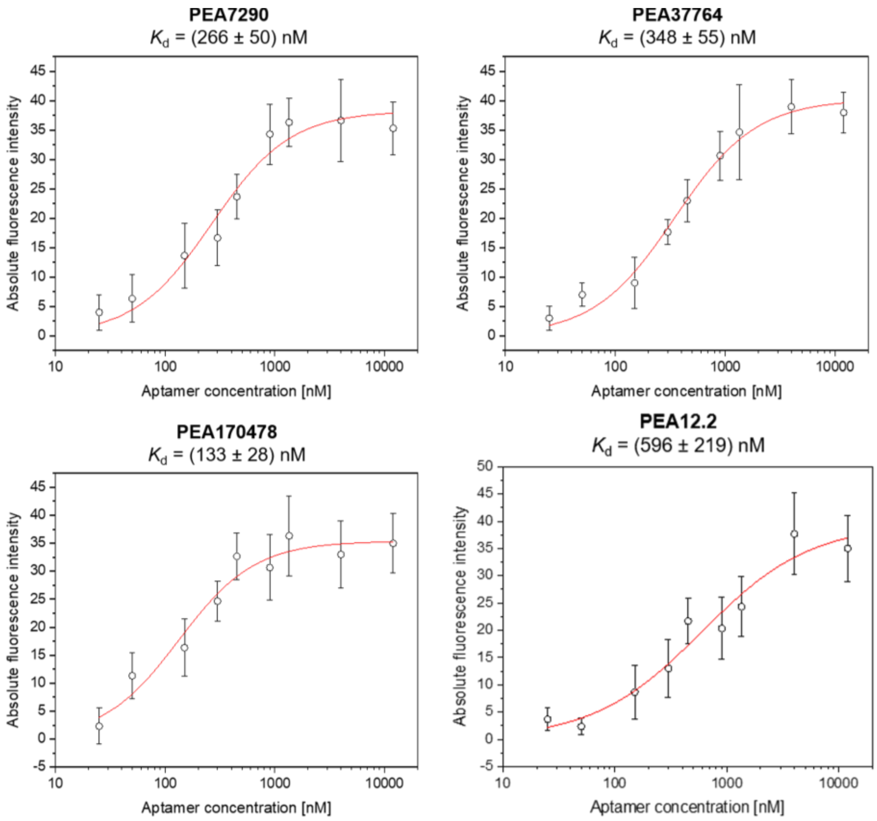


**Figure S9**: Binding curves determined by fluorescence measurement after affinity chromatography for PEA7290, PEA37764, PEA170478, and PEA12.2 with PEA-beads. The plots show the HILL fit of the arithmetic mean with standard deviation of the triplicate measurement at each aptamer concentration. The aptamers were analysed as 90 nt aptamers hybridised to oligos complementary to the primer sequences at 25 °C in PBS + 1 mM $MgCl_2$.

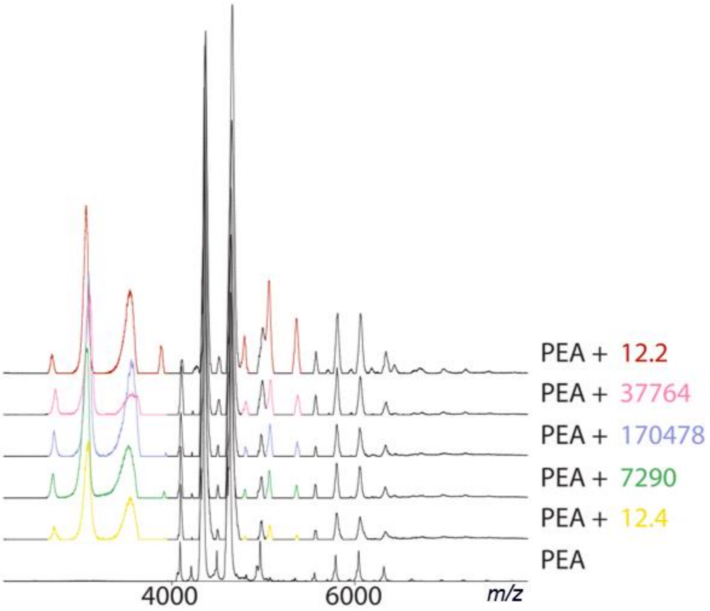


**Figure S10**: Complexes of PEA and selected PEA aptamers observed by nano-ESI-nMS. Coloured peaks of lower *m/z* are from the DNA aptamers whereas coloured peaks between 4000–6000 *m/z* are from the protein-aptamer complex.

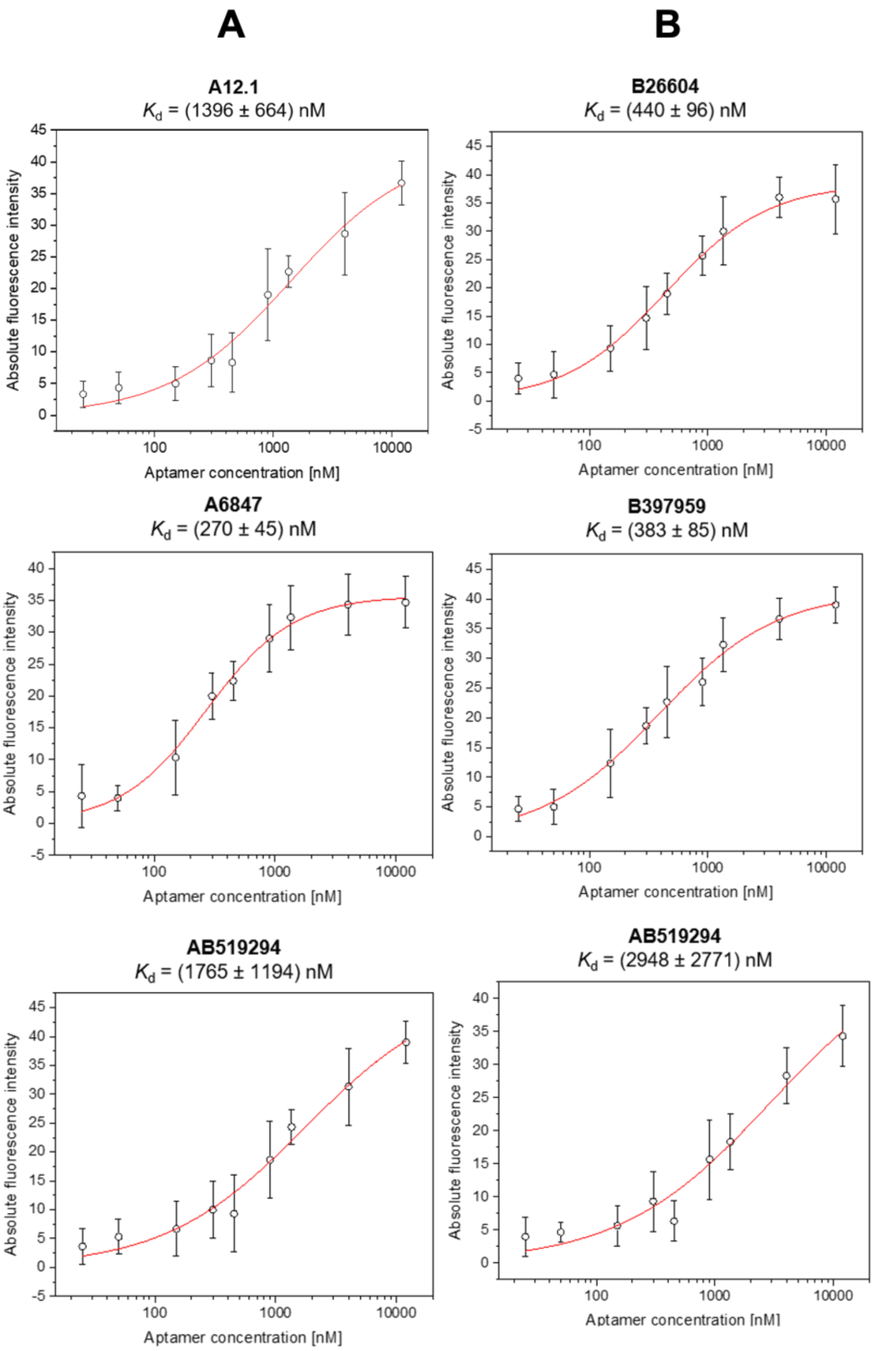


**Figure S11**: Binding curves determined by fluorescence measurement after affinity chromatography for LecA and LecB aptamers with LecA-beads (**A**) or LecB-beads (**B**). The plots show the HILL fit of the arithmetic mean with standard deviation of the triplicate measurement at each aptamer concentration. The aptamers were analysed as 90 nt aptamers hybridised with oligos complementary to their primer sequences in GAMBLES solution at 37 °C.

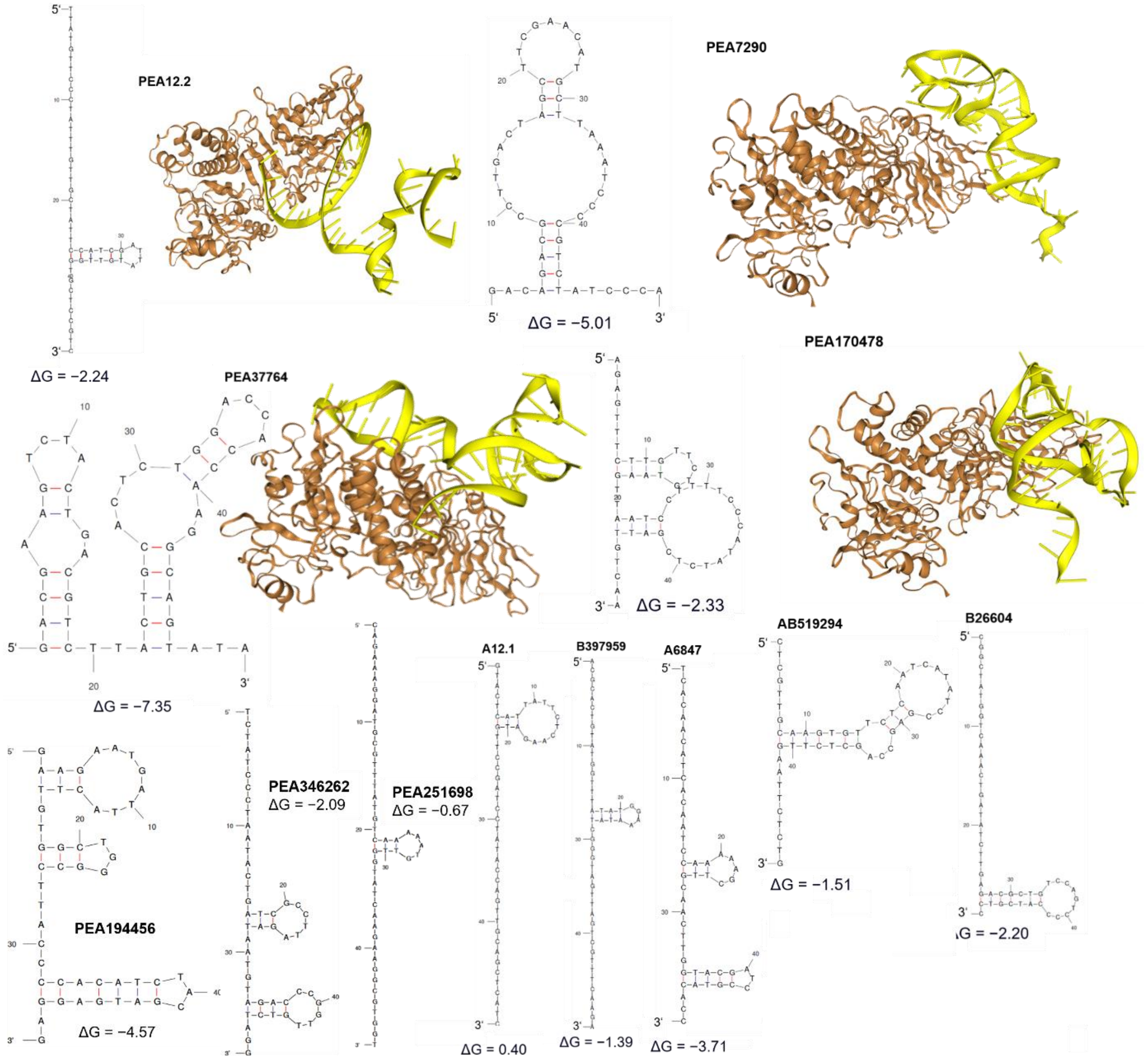


**Figure S12**: Predicted secondary structure and interactions of all aptamers presented in this study. For the best PEA aptamers, additionally molecular docking studies were performed by the HDOCK Server using the DNA’s 3D structure prediction by RNAComposer based on the secondary structure predicted by mfold. The ΔG values are given as [kcal/mol] at 25 °C.

### Supplementary Tables

**Table S1:** Oligonucleotides used, except for the aptamers given in Tab. 1, with their sequence. All oligonucleotides were produced by IDT (Coralville, IA, USA). The manufacturer's designations were used for the modifications: /5AmMC12/: 5' Amino modifier C12; /5IRD700/: 5’ IR-Dye 700; /5Cy5/: 5’ Cy5 dye.

| **Designation** | **Length** | **Sequence (5′–3′)** | **Synthesis scale** |
|---|---|---|---|
| Aptamer library | 90 nt | TCGCACATTCCGCTTCTACC-N50-CGTAAGTCCGTGTGTGCGAA | 2 µmol |
| Forward primer (FP) | 20 nt | TCGCACATTCCGCTTCTACC | 0.1 µmol |
| Complementary FP (FPc) | 20 nt | GGTAGAAGCGGAATGTGCGA | 0.1 µmol |
| Reverse primer (RP) | 20 nt | TTCGCACACACGGACTTACG | 0.1 µmol |
| AminoC12-RP | 20 nt | /5AmMC12/TTCGCACACACGGACTTACG | 25 nmol |
| Fw Primer Seq adaptor | 57 nt | TCGTCGGCAGCGTCAGATGTGTATAAGAGACAGNNNNTCGCACATTCCGCTTCTACC | 25 nmol |
| Rv Primer Seq adaptor | 60 nt | GTCTCGTGGGCTCGGAGATGTGTATAAGAGACAGGAATTCTTCGCACACACGGACTTACG | 25 nmol |
| Fiducial mark (FM) | 50 nt | GTCGTATGCATAAACGAGCCGCACGAACCAGAGAGCATAAAGAGGACCTC | 25 nmol |
| FM Fw Primer Seq adaptor | 57 nt | TCGTCGGCAGCGTCAGATGTGTATAAGAGACAGNNNNGTCGTATGCATAAACGAGCC | 25 nmol |
| FM Rv Primer Seq adaptor | 60 nt | GTCTCGTGGGCTCGGAGATGTGTATAAGAGACAGGAATTCGAGGTCCTCTTTATGCTCTC | 25 nmol |
| FM cy5 complement (FM-A) | 26 nt | /5Cy5/TCGTGCGGCTCGTTTATGCATACGAC | 100 nmol |
| phiX IR700 complement (FM-C) | 20 nt | /5IRD700/GACGTGTGCTCTTCCGATCT | 250 nmol |
| EcoRI cy5 Rv adapter | 20 nt | /5Cy5/TGTATAAGAGACAGGAATTC | 100 nmol |
| PEA R14.33[9] | 80 nt | TGTACCGTCTGAGCGATTCGTACCATAGGGTGCTTTTCAAGGCCACACGTTAGTGTAAGCCAGTCAGTGTTAAGGAGTGC | 100 nmol |
| Fw Primer Seq adaptor for PEA R14.33 | 57 nt | TCGTCGGCAGCGTCAGATGTGTATAAGAGACAGNNNNTGTACCGTCTGAGCGATTCG | 25 nmol |
| Rv Primer Seq adaptor for PEA R14.33 | 60 nt | GTCTCGTGGGCTCGGAGATGTGTATAAGAGACAGGAATTCGCACTCCTTAACACTGACTG | 25 nmol |
| PEA170478 | 50 nt | AGAGTTTCTTGTTCTGTAAGTTAATCCTTTTCCCATATCTCGATTGTCAA | 100 nmol |
| PEA194456 | 50 nt | GAAGAATGATTACTTGTGGCTGGGCCTTACCCCACATCTACGATGAGGAG | 100 nmol |
| PEA346262 | 50 nt | TCTATCCCTAATACTGATCGCCTTAGATAATGTAGACCCGGTTGTCTAGG | 100 nmol |
| PEA251698 | 50 nt | TCAGAAAGGATGCGTTTATGTCAAAAATGTTGGTATCAAGAAGGCGTGGT | 100 nmol |
| PEA7290 | 50 nt | GACAGACGCCTTGACTAGCTTCGAACATGCTTAAATCCCCGTCTATCCCA | 100 nmol |

**Table S2**: Exemplary assignment of the external valve with designations. c: concentration.

| Port | Description | Port | Description |
|---|---|---|---|
| 1 | Air | 7 | Target c3 |
| 2 | NaOH | 8 | Target c4 |
| 3 | Hybridisation buffer | 9 | Target c5 |
| 4 | Pre-imaging buffer | 10 | Target c6 |
| 5 | Target c1 | 11 | Target c7 |
| 6 | Target c2 | 12 | Formamide |

**Table S3**: Exemplary assignment of the custom-filled cartridge with designations. c: concentration, PR2: Illumina proprietary carrier buffer, FM: labelled complementary fiducial mark oligonucleotide, IMT: Illumina proprietary incorporation mix for first base.

| Port | Description | Port | Description |
|---|---|---|---|
| 1 | NaOH | 12 | Target c8 |
| 2 | Hybridisation Buffer | 13 | Target c9 |
| 3 | Incorporation buffer (PR2) | 14 | Target c10 |
| 4 | Pre-imaging buffer | 15 | - |
| 5 | Target c1 | 16 | - |
| 6 | Target c2 | 17 | - |
| 7 | Target c3 | 18 | Primer mix |
| 8 | Target c4 | 19 | FM1 |
| 9 | Target c5 | 20 | FM2 |
| 10 | Target c6 | 21 | IMT |
| 11 | Target c7 | 22 | Formamide |

**Table S4**: Description of the newly created folders of the MiSeq.

| Folder | Description |
|---|---|
| BatchFiles | Contains batch files that are automatically executed at start-up, e. g., for initializing the valve or setting up an alias at system start-up |
| HiTS_FLIP_Inis | Contains ini files to provide the valve assignment |
| HiTS_FLIP_Logs | Contains log files to store all executed actions e. g., switching the valve, detecting a new cycle |
| PythonCode | Location of Python scripts |
| ViciValveLogs | Contains log files of the valve. Each switching is documented here |
| HiTS_FLIP_Recipe | Contains XML for HiTS-FLIP |

**Table S5**: Overview of the additionally installed Python packages with their usage in alphabetical order.

| Package | Usage |
|---|---|
| ftdi-serial | Retrieve serial address |
| matplotlib | Evaluation of the HiTS-FLIP |
| numpy | Evaluation of the HiTS-FLIP |
| os | Environment control |
| pandas | Read out the ini file (.xlsx) |
| pyserial | Communication with valve |
| scipy | Evaluation of the HiTS-FLIP |
| thread | Copying of CIF files on further processor core |
| Tk | Output of error messages |
| vicivalve | Control of the external valve |

**Table S6**: Overview of the rounds of FISHing of the SELEX for LecA.

| SELEX round | Input ssDNA [pmol] | Target/ counter-selection beads volumes [µL] | Incubation time [min] | Number of washing steps | Washing volume [µL] | Counter-selection | Elution |
|---|---|---|---|---|---|---|---|
| 1 | 2000 | 16/16 | 40 | 1 | 200 | yes | Heat |
| 2 | 209 | 16/16 | 40 | 1 | 200 | yes | Heat |
| 3 | 242 | 16/16 | 40 | 1 | 200 | yes | Heat |
| 4 | 85 | 16/16 | 40 | 1 | 200 | yes | Heat |
| 5 | 81 | 16/- | 40 | 2 | 200 | no | Heat |
| 6 | 184 | 16/- | 40 | 2 | 200 | no | Heat |
| 7 | 179 | 16/- | 35 | 2 | 300 | no | Heat |
| 8 | 141 | 16/- | 30 | 3 | 300 | no | Heat |
| 9 | 98 | 16/- | 25 | 3 | 400 | no | Heat |
| 10 | 75 | 16/- | 20 | 3 | 400 | no | Heat |
| 11 | 123 | 16/- | 15 | 3 | 600 | no | Competitive |
| 12 | 82 | 16/- | 15 | 3 | 600 | no | Competitive |

**Table S7**: Overview of the rounds of FISHing of the SELEX for LecB.

| SELEX round | Input ssDNA [pmol] | Target/ counter-selection beads volumes [µL] | Incubation time [min] | Number of washing steps | Washing volume [µL] | Counter-selection | Elution |
|---|---|---|---|---|---|---|---|
| 1 | 2000 | 16/16 | 40 | 1 | 200 | yes | Heat |
| 2 | 266 | 16/16 | 40 | 1 | 200 | yes | Heat |
| 3 | 112 | 16/16 | 40 | 1 | 200 | yes | Heat |
| 4 | 168 | 16/16 | 40 | 1 | 200 | yes | Heat |
| 5 | 140 | 16/- | 40 | 2 | 200 | no | Heat |
| 6 | 99 | 16/- | 40 | 2 | 200 | no | Heat |
| 7 | 168 | 16/- | 35 | 2 | 300 | no | Heat |
| 8 | 88 | 16/- | 30 | 3 | 300 | no | Heat |
| 9 | 96 | 16/- | 25 | 3 | 400 | no | Heat |
| 10 | 84 | 16/- | 20 | 3 | 400 | no | Competitive |
| 11 | 102 | 16/- | 15 | 3 | 600 | no | Heat |
| 12 | 99 | 16/- | 15 | 3 | 600 | no | Heat |

**Table S8**: Overview of the rounds of FISHing of the SELEX for PEA.

| SELEX round | Input ssDNA [pmol] | Target/ counter-selection beads volumes [µL] | Incubation time [min] | Number of washing steps | Washing volume [µL] | Counter-selection | Elution |
|---|---|---|---|---|---|---|---|
| 1 | 2000 | 16/16 | 40 | 1 | 200 | yes | Heat |
| 2 | 345 | 16/16 | 40 | 1 | 200 | yes | Heat |
| 3 | 185 | 16/16 | 40 | 1 | 200 | yes | Heat |
| 4 | 178 | 16/16 | 40 | 1 | 200 | yes | Heat |
| 5 | 111 | 16/- | 40 | 2 | 200 | no | Heat |
| 6 | 163 | 16/- | 40 | 2 | 200 | no | Heat |
| 7 | 140 | 16/- | 35 | 2 | 300 | no | Heat |
| 8 | 138 | 16/- | 30 | 3 | 300 | no | Heat |
| 9 | 100 | 16/- | 25 | 3 | 400 | no | Heat |
| 10 | 117 | 16/- | 20 | 3 | 400 | no | Competitive |
| 11 | 97 | 16/- | 15 | 3 | 600 | no | Competitive |
| 12 | 101 | 16/- | 15 | 3 | 600 | no | Competitive |

**Table S9**: Parameters of nano-ESI-nMS.

| **Device** | | Modified Micromass qToF |
|---|---|---|
| **Source** | Gas | Argon |
| | Capillary | 1.35 kV |
| | Cone | 150 V |
| | Extractor | 0 V |
| | RF Lens | 0.1 V |
| | Multiplier | 550 V |
| | MCP | 1900 V |
| | Temperature | 80 °C |
| | Desolvation Temp | 20 °C |
| **MS** | LM Res | 10 |
| | HM Res | 10 |
| | Pressure in Collision Cell | 1.3–1.5 kV × $10^{-2}$ mbar |
| | Collision | 20–180 V |
| | Ion Energy | 1 |
| | Steering | 0 |
| | Entrance | 70 |
| | Pre-filter | 5 |
| | Multiplier | 550 |
| | MCP | 1900 V |
